# WS6 enhances expansion of primary human airway basal cells and supports clonal genome editing

**DOI:** 10.1101/2025.09.15.675861

**Authors:** Jessica C. Orr, Elizabeth K. Haughey, Andrew S. Farr, David R. Pearce, Niamh A. McCarthy, Shalini Kamu Reddy, Maral J. Rouhani, Charlotte Percival, Isabelle Rose, Anna Straatman-Iwanowska, Rebecca B. Dale, Aleksandra Z. Szejn, Megan Guthrie, Giada Benedetti, Buthainah M. Ahmed, Olivia R. Pape, Juan Moisés Ocampo-Godinez, Elizabeth F. Maughan, Colin R. Butler, Dale A. Moulding, Alexandra Y. Kreins, Giovanni Giuseppe Giobbe, Paolo De Coppi, William Grey, Antonella F. M. Dost, Robert A. Hirst, Deborah L. Baines, Yuki Ishii, Christopher O’Callaghan, Sam M. Janes, Robert E. Hynds

## Abstract

Fundamental airway epithelial biology, disease modeling and regenerative medicine research rely on the ability to expand and manipulate primary human airway basal cells, often from small clinical samples. Basal stem/progenitor cells from (pseudo)stratified epithelia can be expanded in 3T3-J2 fibroblast feeder cell co-culture systems, and the addition of the ROCK inhibitor Y-27632 enhances proliferation and culture longevity, a phenomenon that has been described as ‘conditional reprogramming’. Here, we developed ‘EpMED’, a feeder-based FAD medium supplemented with Y-27632 and WS6. Evaluation of nasal airway basal cells obtained by brush biopsy showed that inclusion of WS6 increased colony formation, cumulative expansion and culture duration in this system. Evidence from colony formation assays using bronchial, small airway, skin and thymic cultures was suggestive of wider relevance to other epithelia. Nasal basal cells maintained in EpMED retained basal cell marker expression, clonogenicity, multilineage differentiation, epithelial barrier and ion transport functions. The cells had no detectable copy number alterations and showed no evidence of tumor formation after subcutaneous xenotransplantation. EpMED-expanded nasal basal cells engrafted and regenerated differentiated epithelium in a tracheal injury model in immunodeficient mice, and were capable of serial transplantation. In a second application, we generated clonal CRISPR-Cas9 genome edited nasal cultures, introducing *DNAH5*- or *DNAI2*-knockout to model primary ciliary dyskinesia. These findings establish EpMED as a scalable platform for expanding primary human airway basal cells from low-input samples and for generating clonal airway disease models.

## Introduction

Robust *in vitro* cell culture systems for primary human epithelial cells underpin key applications in disease modeling, functional genomics, and regenerative medicine. The discovery that mouse embryonic fibroblasts can act as feeder cells enabled primary epithelial cell culture for the first time (Rheinwald & Green, 1975). The initial medium, which contained fetal calf serum and hydrocortisone, was later refined to additionally include adenine, cholera toxin, epidermal growth factor, insulin, and triiodothyronine (‘FAD medium’; reviewed in (Hynds, Bonfanti, et al., 2018)). This formulation supports the expansion of epithelial basal stem/progenitor cells from multiple tissues, including skin, cornea, esophagus, thymus, and respiratory tract (Adamo et al., 2025; Compton et al., 1998; Pellegrini, Golisano, et al., 1999; Ragazzini et al., 2023; Vietri Rudan et al., 2020). The addition of the Rho kinase inhibitor Y-27632 to FAD enhances epithelial cell colony formation potential, proliferation, and longevity (Butler et al., 2016; Chapman et al., 2010; Liu et al., 2012; Terunuma et al., 2010).

Despite improvements to the 2D culture of epithelial cells over recent decades, the ability to extensively expand primary human cells from clinically available biopsy samples remains challenging, particularly from patients with epithelial diseases that limit stem/progenitor cell function (Ghosh et al., 2018; Hong et al., 2022). Here, to focus on expansion of cells from low-input clinical material, we use nasal brush biopsies as the primary cell source. Nasal brush biopsies are more readily available than other biopsy types, they provide a minimally invasive source of airway basal cells but yield relatively few starting cells compared to surgical samples, making them a stringent test of culture robustness.

A previous report described maintaining mouse totipotent stem cells with a small-molecule cocktail (Hu et al., 2023). The cocktail included 1-azakenpaullone, a Wnt pathway activator that we previously found to promote airway basal cell proliferation (Ishii et al., 2025). It also contained TTNPB, a retinoic acid analogue, and WS6, which is reported to bind to proliferation-associated protein 2G4 (PA2G4, or EBP1) and IKKε (Shen et al., 2013), a regulator of the NFkB pathway. Here, we investigate the potential of these small molecules to enhance epithelial cell culture. We demonstrate that the addition of WS6, in combination with Y-27632, to FAD medium enhances colony formation, proliferation, and culture lifespan of primary airway basal cell cultures. We then apply this system to airway transplantation and CRISPR-Cas9 disease modeling, where we develop isogenic, clonal knockout cultures capable of multi-lineage differentiation for *DNAH5* or *DNAI2*, genes in which pathogenic variants are known to cause primary ciliary dyskinesia.

## Results

### WS6 enhances colony formation and proliferation in primary airway cell cultures

To explore the effects of the small molecule cocktail described by Hu et al. (Hu et al., 2023) on human epithelial cells, we first assessed the effects of 1-azakenpaullone, TTNPB, and WS6 on proliferation in primary nasal and bronchial airway cell cultures. WS6 alone, or in combination with the other compounds, increased cell numbers (**Fig. 1A**). Concentration optimization of Y-27632 and WS6 was then performed in bronchial and skin cultures, with the combination of 1-10 μM Y-27632 and 0.1-1 μM WS6 proving most effective. At higher WS6 concentrations (>1 μM), toxicity was observed (**Fig. 1B**).

Colony formation assays provide a functional readout of basal stem/progenitor cell potential and Y-27632 enhances colony formation (Reynolds et al., 2016; Zhou et al., 2013). To assess the effect of WS6, WS6 (100 nM) alone or in combination with Y-27632 (5 μM) was added at the time of assay seeding to nasal basal cells that had been established on 3T3-J2 feeder layers in FAD medium. Alone, WS6 (FAD+WS6) produced a similar effect to Y-27632 (FAD+Y), while the combination (FAD+Y+WS6) further increased colony formation (**Fig. 1C, left**). Prompted by these findings, we next used colony formation assays to assess the activity of WS6 across selected primary epithelial cell sources. Compared to FAD alone, the combination of Y-27632 and WS6 significantly increased colony formation of nasal, bronchial and small airway cells, with a similar trend observed for epidermal and thymic epithelial cells (**Fig. 1C**). We also tested whether WS6 could enhance the initial culture colony forming efficiency of epithelial cells directly from primary human tissue. We dissociated tracheal and nasal biopsies into single cells and sorted EPCAM⁺ cells individually onto 3T3-J2 feeder layers. In all donors tested, WS6 increased the number of cells capable of forming colonies at explant (**Fig. 1D**).

**Figure 1:**
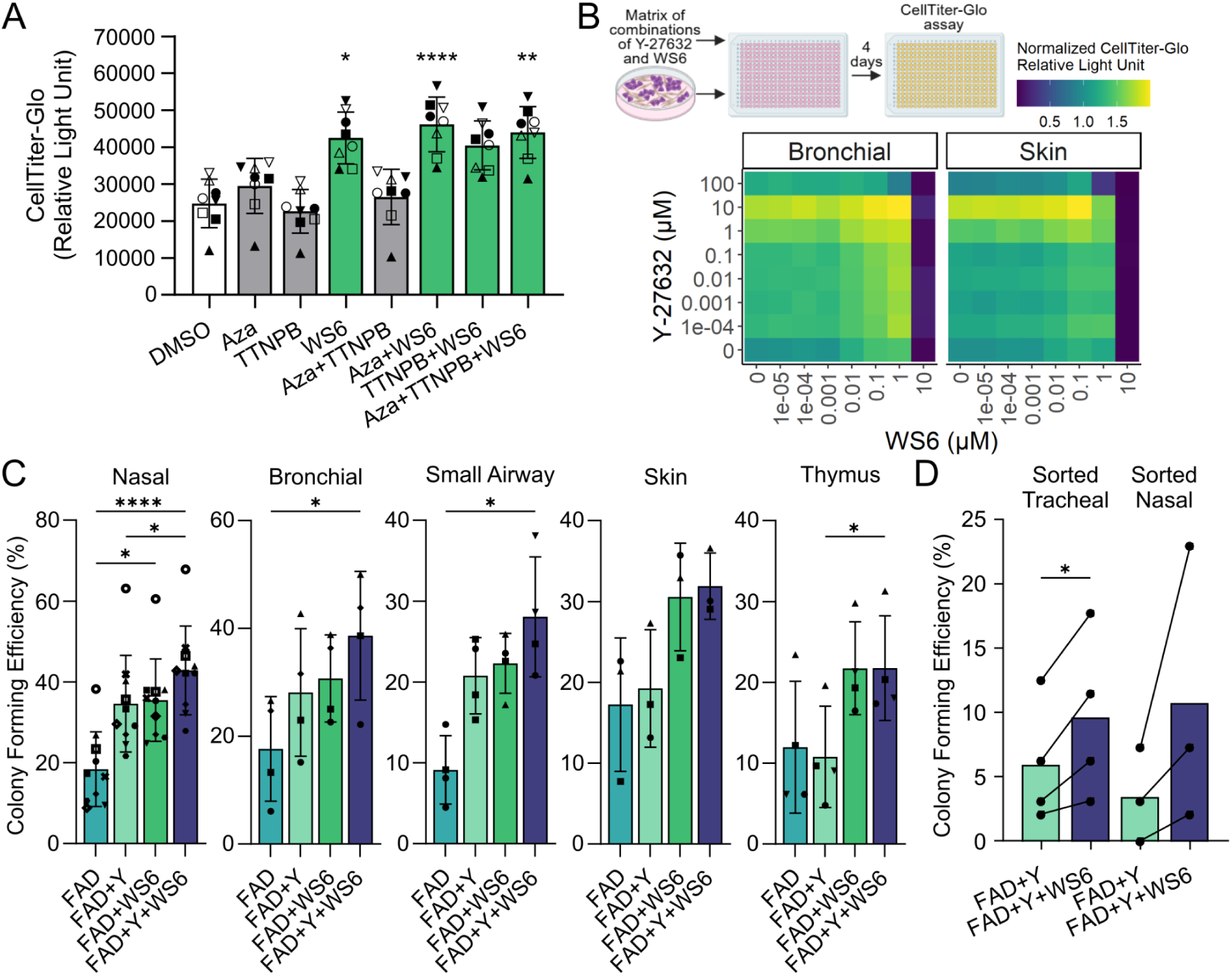
WS6 promotes epithelial basal cell proliferation in 3T3-J2 co-culture. A) Six-day CellTiter-Glo assay to examine the effect of combinations of the small molecules (2.5 µM 1-azakenpaullone (Aza), 0.2 µM TTNPB and 0.5 µM WS6) identified to induce totipotency in mouse embryonic stem cells (Hu et al., 2023) on epithelial basal cell proliferation (n = 4 primary human nasal basal cell donors (filled points) and n = 4 primary human bronchial cell donors (open points), indicated by shapes). Friedman test with Dunn’s multiple comparison test vs DMSO control wells. B) Four-day CellTiter-Glo assays to assess the effect of varying concentrations of Y-27632 and WS6 on cell proliferation in passage three bronchial (left, mean of five donors is shown) and skin (right, mean of two independent cultures from one donor) basal cells previously grown in FAD medium. C) Colony formation assays to assess the effect of Y-27632 (Y), WS6, and the combination of both on colony forming potential of primary human epithelial cells from different tissues (n = 9 nasal, n = 4 bronchial, n = 4 small airway, n = 3 skin, n = 4 thymus donors, indicated by shapes). Friedman test with Dunn’s multiple comparison test. D) Colony formation assays to assess the effect of Y, and the combination of Y and WS6 on colony forming potential of FACS-sorted EPCAM^+^ human tracheal (n = 4 donors) or nasal (n = 3 donors) epithelial cells directly from biopsies. Paired t-test per sample location.

To determine whether the pro-proliferative effect of WS6 extended beyond primary epithelial cells, we first assessed commonly used epithelial cell lines and primary lung fibroblasts. WS6 increased proliferation of *BMI1*-transduced primary nasal cell cultures, but not of A549 cells (a lung adenocarcinoma cell line), H520 cells (a lung squamous cell carcinoma cell line), HBEC3-KT cells (Ramirez et al., 2004) (an airway basal cell line immortalized by transduction with cyclin-dependent kinase 4, CDK4, and human telomerase reverse transcriptase, hTERT), or primary lung fibroblasts (**Supplementary Fig. 1**).

We next asked whether WS6 influences primary human cell culture systems beyond 2D stratified epithelia. 1 μM WS6 increased proliferation in small airway organoids, reduced proliferation of alveolar organoids and had no effect on gastric organoids (**Supplementary Fig. 2**). In CD34+ hematopoietic cell cultures, WS6 reduced the total nucleated cell output while modestly enriching the hematopoietic stem cell (HSC) fraction (**Supplementary Fig. 3A-G**). Colony forming unit (CFU) assays showed that WS6 did not alter hematopoietic progenitor lineage output, with similar proportions of CFU-macrophage (CFU-M), CFU-granulocyte (CFU-G), CFU-granulocyte macrophage (CFU-GM), and CFU-granulocyte erythrocyte macrophage megakaryocyte (CFU-GEMM) colonies across all conditions (**Supplementary Fig. 3H**). Taken together, these findings show that WS6 stimulation of proliferation in human cell cultures is dependent on cell type, tissue origin and culture context and does not represent a universal epithelial proliferation stimulus.

### Long-term expansion of primary nasal airway basal cells in WS6-containing medium

To assess long-term cell expansion in WS6-containing medium, cells from individual nasal brush biopsies were divided between three conditions and cultured in parallel: FAD medium alone, FAD+Y, or FAD+Y+WS6. Since studies converge on cell counts of approximately 1 × 10^6^ cells per nasal brushing and approximately one third of nasal epithelial cells are basal cells, each of three parallel cultures was initiated from approximately 1 × 10^5^ basal cells (Fawcett et al., 2023; Stokes et al., 2014). All three conditions supported early passage expansion of cells with characteristic epithelial cell cobblestone morphology (**Fig. 2A**). Across all conditions, cells maintained cytoplasmic expression of keratin 5 (KRT5), a basal cell intermediate filament protein (Fuchs & Blau, 2020), and nuclear expression of TP63, a transcription factor central to epithelial basal cell identity (Li et al., 2023; Pellegrini et al., 2001) (**Fig. 2B and Supplementary Fig. 4**). Over time, however, cells in FAD alone became enlarged, and developed bright, refractile colony centers consistent with the onset of stratification; this was delayed in FAD+Y conditions and further delayed in FAD+Y+WS6 conditions (**Supplementary Fig. 5**). Cells in FAD+Y+WS6 expanded more rapidly (**Fig. 2C-D**) and could be maintained for extended culture durations (**Fig. 2E and Supplementary Fig. 5 and 6A**), with one donor culture exceeding one year of continuous cell culture and accumulating over 100 passages (**Supplementary Fig. 6A**). This resulted in a greater cell yield in FAD+Y+WS6 than FAD culture conditions from small clinical samples (**Fig. 2F**). When benchmarked against a commercially available feeder-free expansion medium (PneumaCult™-NGEx), FAD+Y+WS6 cultures (on 3T3-J2 feeder layers) were similar at the first passage (**Supplementary Fig. 7A-C**) but subsequently expanded more rapidly and to an overall higher population doubling (**Supplementary Fig. 7A, 7D-E**).

**Figure 2:**
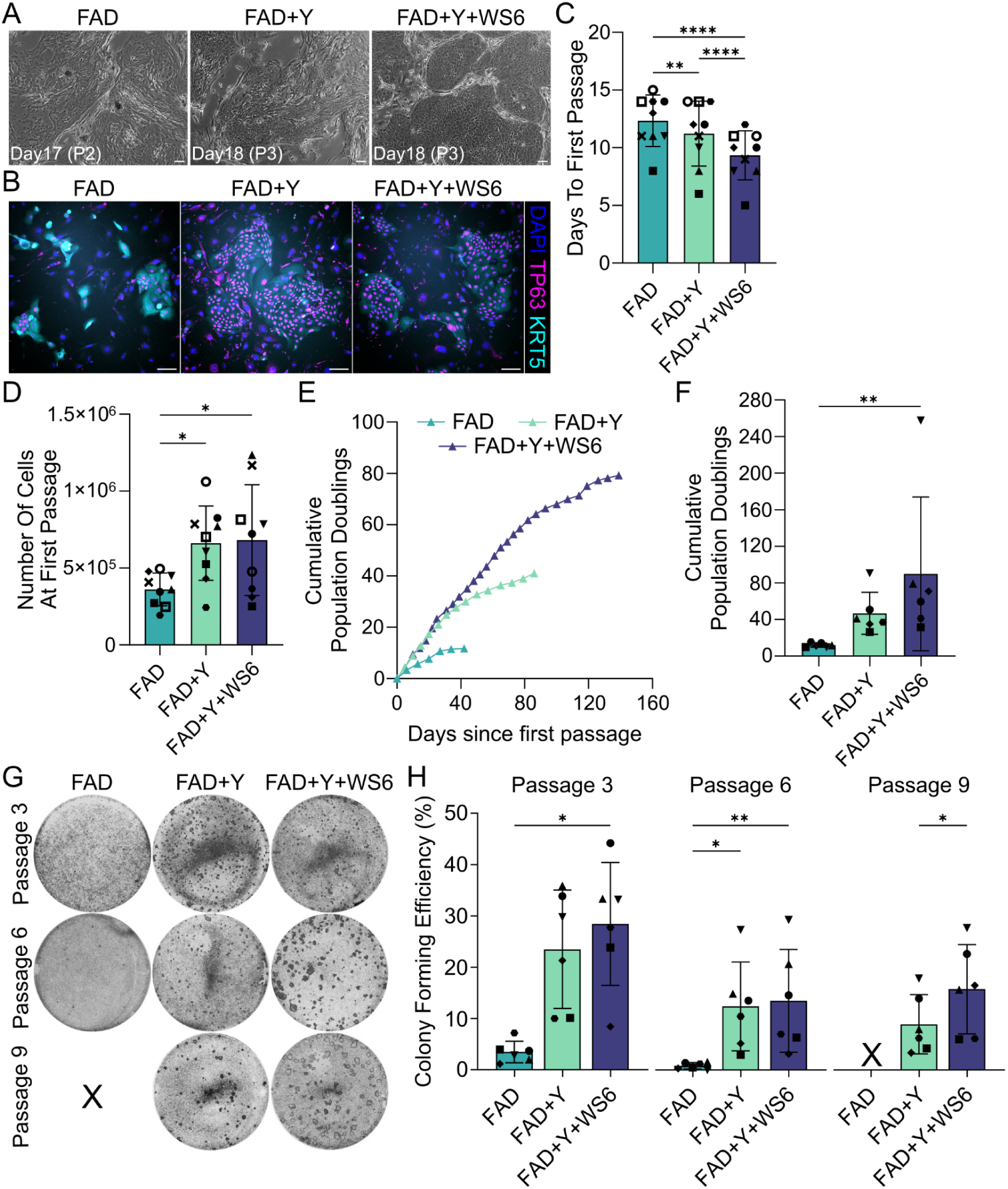
WS6 extends the culture duration of nasal basal epithelial cells. A) Phase-contrast images of nasal epithelial basal cells cultured in FAD (left), FAD+Y-27632 (FAD+Y, center) or FAD+Y+WS6 (right) at indicated days since explant and passages. Scale bar = 100 µm. B) Immunofluorescence staining of passage three nasal epithelial basal cells fixed at day three of culture (TP63, magenta; KRT5, cyan; DAPI, blue). Scale bar = 100 µm. A-B) Images shown from a single donor (n = 6 donors tested). C-D) Initial passaging of donor-matched nasal epithelial basal cells from a single brush biopsy split into three 24-well format wells for culture in FAD, FAD+Y or FAD+Y+WS6. Repeated measures one-way ANOVA with Tukey’s multiple comparison and donor as a matched variable (n = 9 donors, indicated by shape). Days (C) and number of cells collected (D). E) Representative population doublings of nasal epithelial basal cells cultured in FAD, FAD+Y or FAD+Y+WS6 from a single donor (N068N). F) Cumulative population doublings for donor-matched nasal epithelial cells cultured in FAD, FAD+Y or FAD+Y+WS6. Maximum population doublings reported except for the single donor in FAD+Y+WS6 condition that exceeded 100 passages where data represents cumulative population doublings at day 366. Friedman test with Dunn’s multiple comparison (n = 6 donors). G-H) Colony formation assays for donor-matched nasal epithelial basal cells cultured in FAD, FAD+Y or FAD+Y+WS6 at passages three, six, and nine. Representative images from a single donor are shown (G) and colony number quantified (H; n = 6 donors, passages three and six = one-way ANOVA with Tukey’s multiple comparison, passage nine = paired t-test). X indicates that FAD cultures had undergone replicative senescence prior to passage nine. Shapes indicate donors.

Colony formation was consistently higher across passages in FAD+Y and FAD+Y+WS6 cultures than in FAD alone (**Fig. 2G-H**), with the highest colony forming efficiencies seen at early passage in all media (**Supplementary Fig. 6B**). At higher population doublings, more colonies were formed by FAD+Y+WS6 cultures than FAD+Y cultures (**Fig. 2H**), consistent with the preservation of clonogenic potential in FAD+Y+WS6.

In Green’s FAD, the retention of epithelial stem cells is dependent on the presence of 3T3-J2 feeder cells and this remains true with Y-27632 addition (Liu et al., 2012). The addition of WS6 did not eliminate the need for feeder cells to support long-term culture (**Supplementary Fig. 8A-C**). In FAD+Y+WS6 feeder cell withdrawal led to reduced culture longevity and fewer total population doublings (**Supplementary Fig. 8A-B**), and did not support maintenance of epithelial cobblestone morphology (**Supplementary Fig. 8C**) in nasal basal cells from five donors.

Previous studies have shown improved lentiviral transduction of epithelial basal cells with Y-27632 (Horani et al., 2013; van den Bogaard et al., 2012). While Y-27632 improved epithelial cell transduction with fluorescent reporter vectors in our studies as expected, WS6 conferred no benefit, alone or in combination with Y-27632 (**Supplementary Fig. 9**).

Low-pass whole-genome sequencing (0.4x coverage) performed on nasal airway basal cells from seven donors after 9-10 passages showed stable copy number profiles without detectable copy number alterations (**Fig. 3A**). We also transplanted cells from six donors, expanded for 12-14 passages in FAD+Y+WS6, into NOD *scid* gamma (NSG) mice subcutaneously in Matrigel. No tumors were observed in the six mice during the 12-week observation period. In 3/6 mice, we were able to recover the injected Matrigel which contained epithelial structures resembling differentiated airway tissue, consistent with the retention of airway differentiation capacity *in vivo* (**Fig. 3B**).

**Figure 3:**
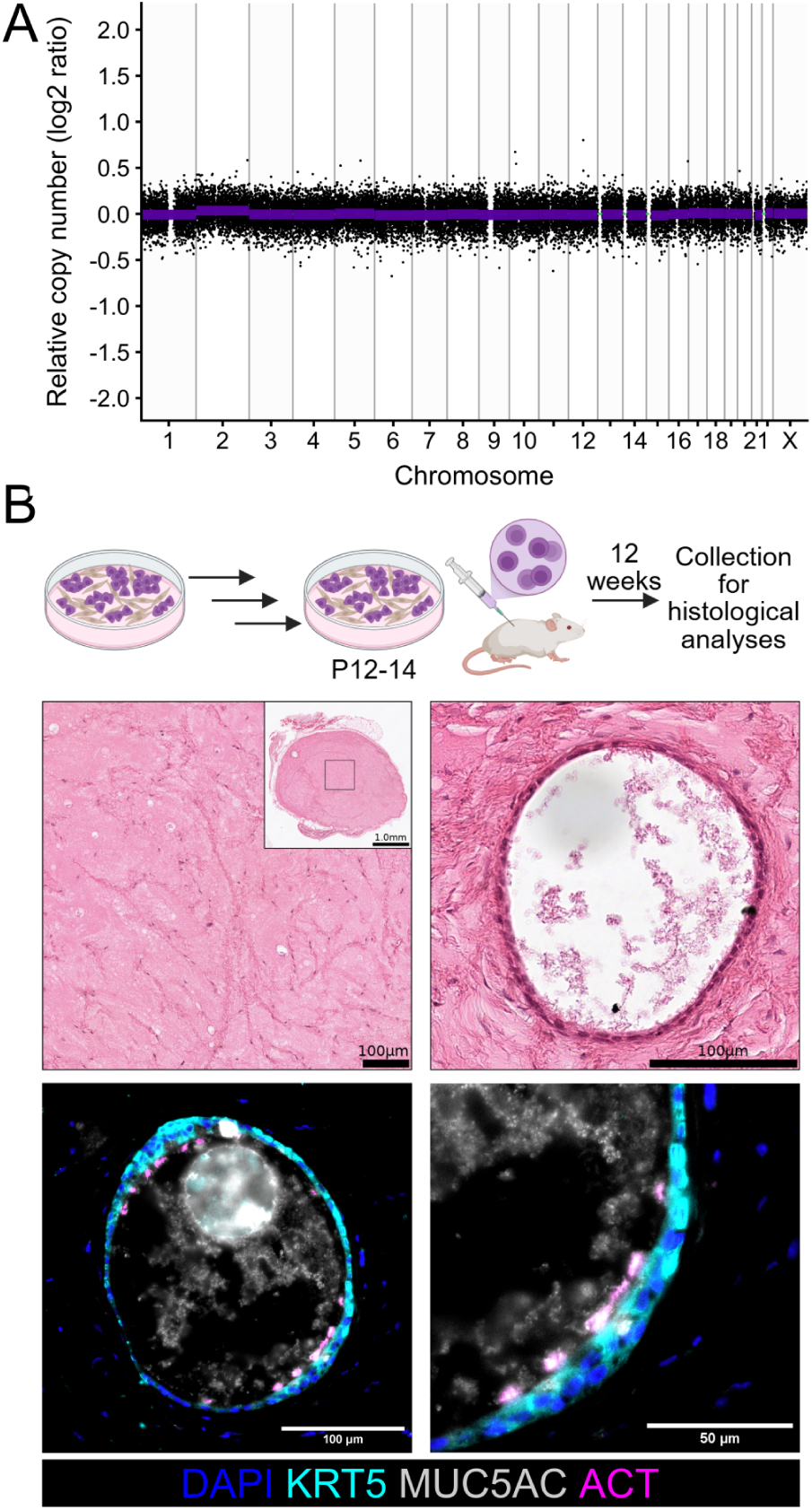
Cells cultured in WS6 had no detectable copy number alterations and did not form tumors in mice. A) A representative copy number profile from low-pass whole-genome sequencing of nasal epithelial cells expanded in FAD+Y-27632(Y)+WS6 (n = 6 donors at passage nine, n = 1 donor at passage ten, N039 shown). Points indicate unflagged bins and purple horizontal lines indicate copy number segments. B) Subcutaneous injection of nasal epithelial basal cells expanded in FAD+Y+WS6 for 12-14 passages. Tumors formed in 0/6 mice (1 unique donor/mouse). Hematoxylin and eosin staining (upper) and immunofluorescence staining (lower; KRT5, cyan; MUC5AC, grey; acetylated tubulin, ACT, magenta; DAPI, blue) representative of 3/6 mice in which epithelial structures could be found (N035, N036, N045) within the extracellular matrix recovered 12 wk after injection, data for N036 shown. Scale bar lengths are indicated within the figure panel.

For *in vitro* and *in vivo* applications, epithelial basal cells must retain the capacity for tissue-appropriate differentiation. When induced to differentiate in air-liquid interface (ALI) cultures after four passages in FAD medium, FAD+Y or FAD+Y+WS6 (**Fig. 4A**), nasal epithelial cells formed a pseudostratified epithelium containing both mucosecretory and ciliated cells (**Fig. 4B-C and Supplementary Fig. 10A**). At passages 4 or 10, FAD+Y and FAD+Y+WS6 cultures produced comparable proportions of ciliated cells (**Fig. 4D**) but for both 2D culture conditions the passage 10 ALI cultures produced a higher proportion of MUC5AC⁺ mucosecretory cells than the passage 4 ALI cultures (**Fig. 4D**). Cells from FAD+Y+WS6 cultures were able to differentiate in 3D organoid assays throughout long-term culture (**Supplementary Fig. 11A-C**), with one donor retaining full differentiation capacity after 38 passages and mucosecretory, but not ciliated, cell differentiation after 100 passages (**Supplementary Fig. 11D**). These findings support the retention of multilineage differentiation capacity in nasal airway basal cells even after extended culture in FAD+Y+WS6.

We next investigated cellular functionality in ALI cultures. First, across all passages and medium compositions, there were no significant differences in the transepithelial electrical resistance (TEER) measurements, suggesting maintained epithelial integrity (**Supplementary Fig. 10B**). Second, the ciliary beat frequency of ciliated cells was within the expected normal range (Jorissen & Bessems, 1995; Smith et al., 2014) (**Fig. 4E and Supplementary Fig. 10C**). Area covered by motile cilia was increased in FAD+Y and FAD+Y+WS6 ALI cultures relative to FAD (**Fig. 4E and Supplementary Fig. 10C**). Third, ion channel function was preserved in FAD+Y+WS6 ALI cultures, which responded appropriately to pharmacological agents across multiple cell passages: amiloride, indicating activity of the amiloride-sensitive epithelial Na⁺ channel (ENaC); forskolin and CFTRinh-172, indicating cAMP-regulated CFTR activity; and ATP, indicating calcium-activated Cl⁻ channel (CaCC) activity. Ouabain inhibited all ion channel activity, consistent with Na⁺/K⁺-ATPase activity (**Fig. 4F and Supplementary Fig. 12**).

**Figure 4:**
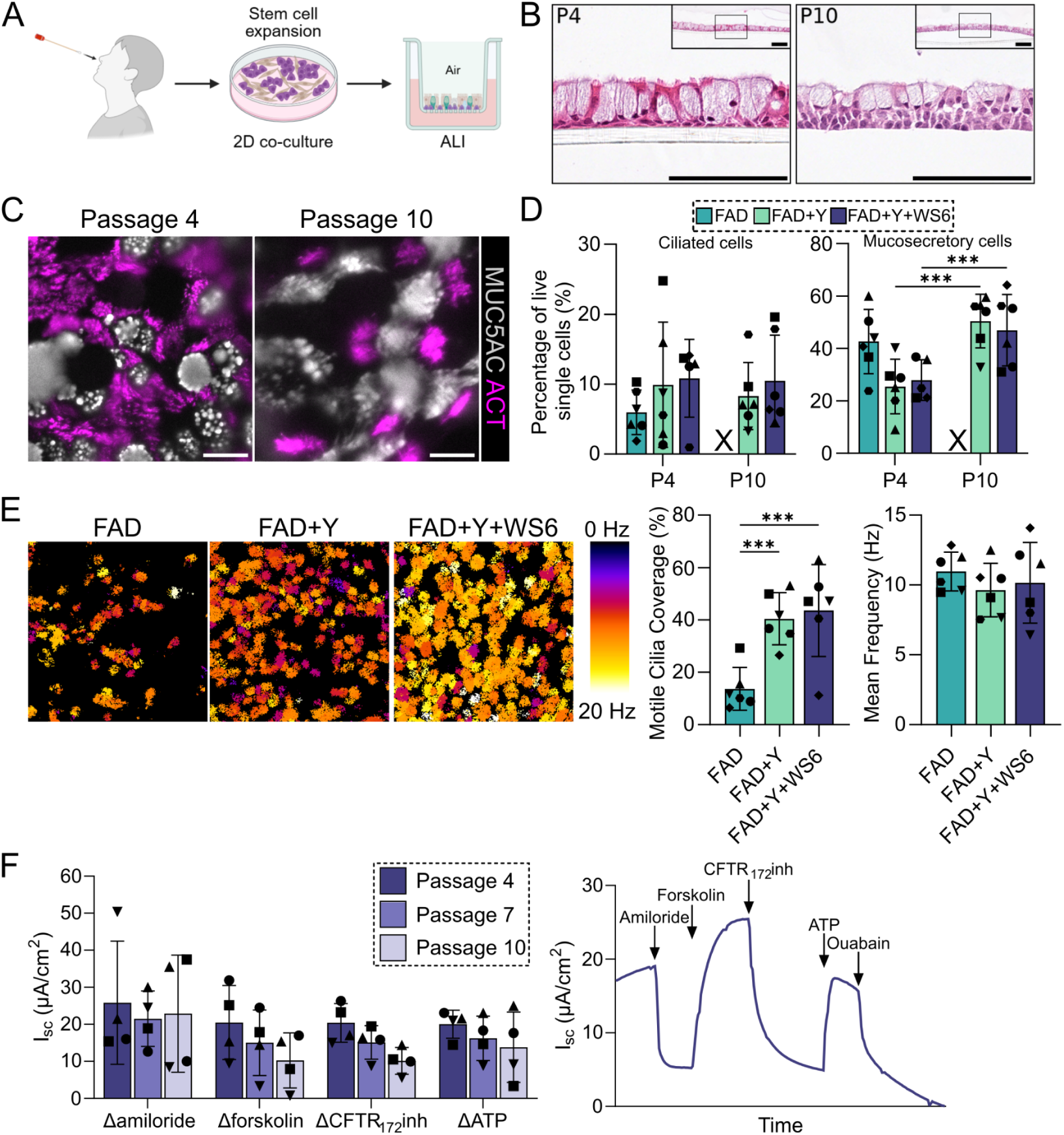
Nasal epithelial cells retain the capacity for multipotent differentiation during extended culture. A) Schematic diagram of nasal epithelial basal cell differentiation in air-liquid interface (ALI) cultures. B) Hematoxylin and eosin staining of formalin-fixed, paraffin-embedded ALI cultures from passage four (P4, left) and P10 (right) cells previously expanded in FAD+Y-27632(Y)+WS6. Scale bar = 100 µm C) Whole-mount immunofluorescence staining of ALI cultures from P4 (left) and P10 (right) cells previously cultured in FAD+Y+WS6 (acetylated tubulin, ACT, magenta; MUC5AC, grey). Scale bar = 20 µm. D) Proportions of epithelial cell types from P4 and P10 ALI cultures of nasal epithelial basal cells from flow cytometry. X indicates that FAD cultures had failed prior to P10 and thus could not be used for ALI experiments (n = 5-6 donors, Two-way ANOVA with Tukey’s multiple comparison per cell type). E) Ciliary beat frequency analysis of differentiated ALI cultures from P4 cells previously expanded in FAD, FAD+Y or FAD+Y+WS6. Coverage and beat frequency of moving cilia are displayed for representative fields of view of FAD, FAD+Y and FAD+Y+WS6 conditions from a single donor (N066N, left). Mean motile cilia coverage and cilia beat frequency is quantified (right). Each data point represents the average frequency of all moving cilia from five fields of view in three technical replicate wells (n = 6 donors; indicated by shape). One-way ANOVA with Tukey’s multiple comparison. F) Ussing chamber analysis of ion transport in ALI cultures derived from FAD+Y+WS6 cultures at passages four, seven and ten. Change in short-circuit current (I_sc_) of day 30 ALI cultures in response to amiloride (100 µM, apical), forskolin (10 µM, bilateral), CFTRinh-172 (10 µM, apical) and ATP (100 µM, apical), points represent the mean of quadruplicate samples for each individual donor, N = 4 donors; indicated by shape (left). Two-way ANOVA performed. Representative I_sc_ trace from a single donor (N067N) at passage 4 (right).

### In vivo transplantation of EpMED-expanded nasal airway basal cells

To assess *in vivo* regenerative potential, we transplanted human nasal basal cells into the tracheas of immunodeficient NSG mice. Cells were isolated and expanded in FAD+Y+WS6 and transduced with a ZsGreen-luciferase lentiviral vector for tracking. Tracheas were preconditioned by oropharyngeal polidocanol instillation, a detergent that denudes the epithelium (Borthwick et al., 2001), before delivery of cells 5 hours later (Ma et al., 2023) (**Fig. 5A**). Engraftment was detected in all 11 recipients by In Vivo Imaging System (IVIS) imaging at day 7, though signal intensity varied between recipients (**Fig. 5B**). Luciferase activity persisted throughout the 28-day experiment, with a gradual reduction over time (**Fig. 5B**), and was localized specifically to the trachea in post-mortem imaging (**Fig. 5C**). At the end-point, whole-mount (**Fig. 5D**) and histological analyses (**Fig. 5E**) confirmed engraftment of human basal cells and their differentiation to mucosecretory and ciliated cells *in vivo* (**Fig. 5E**).

**Figure 5:**
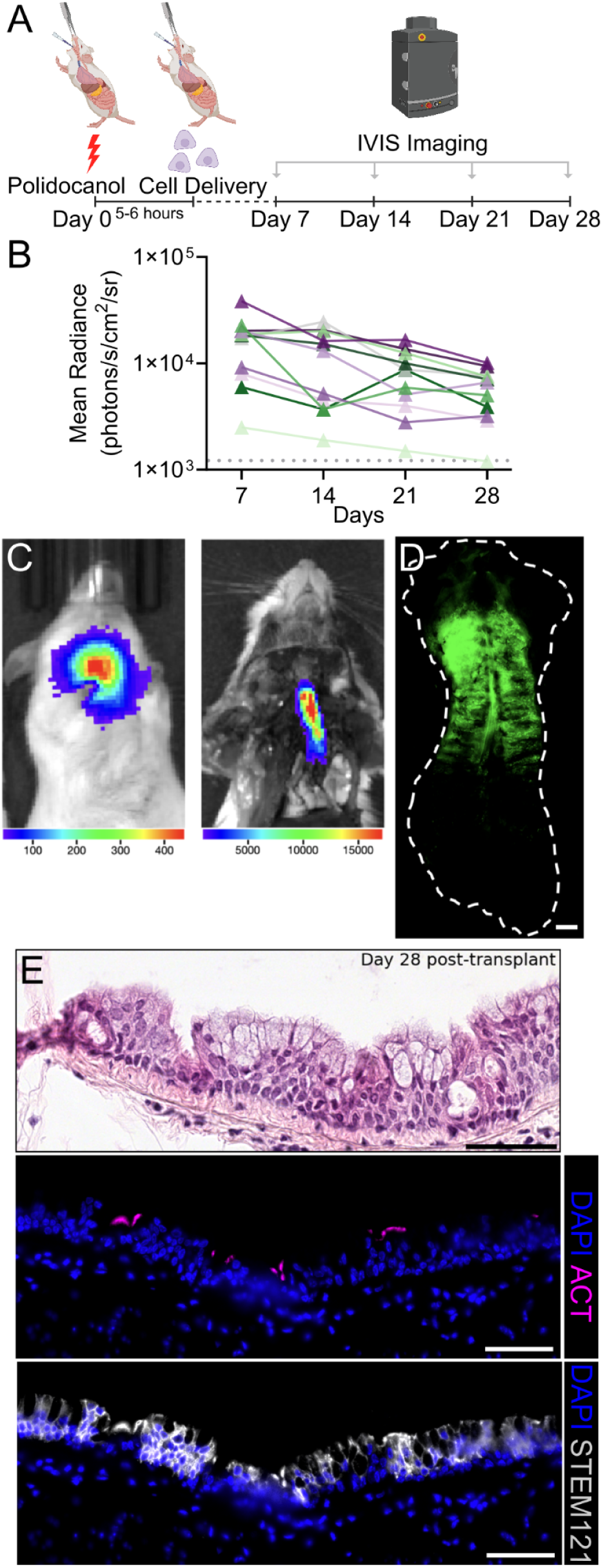
Tracheal transplantation of human basal cells expanded in FAD+Y+WS6. A) Schematic diagram of ZsGreen-luciferase transduced nasal epithelial basal cell transplantation in the trachea of immunodeficient NSG mice and monitoring by luciferase bioluminescence imaging (IVIS). B) Quantification of IVIS signal over 28 d following cell engraftment (n = 11 mice). Dotted line indicates background signal. C) IVIS imaging of a living mouse 7 d after transplantation with human basal cells (left) and the same mouse post-mortem (right), demonstrating the tracheal specificity of cell engraftment. D) Whole-mount imaging of ZsGreen in a freshly excised trachea 28 d following basal cell transplantation. Scale bar = 1 mm. E) Hematoxylin and eosin staining of sections from a mouse trachea 28 d after basal cell transplantation (upper) and immunofluorescence staining demonstrating ciliated cell differentiation (acetylated tubulin, ACT, ciliated cells, magenta) and human origin (STEM121, grey). Scale bar = 50 µm.

Serial transplantation provides a functional test of retained self-renewal and multilineage regenerative potential (Daniel et al., 1968). In a separate transplantation experiment, donor-matched human nasal basal cells expanded in FAD+Y+WS6 and transduced with ZsGreen-Luciferase were transplanted into five preconditioned mice (**Fig. 6A**). Engraftment was detected by IVIS imaging in 4/5 mice across six weeks (**Fig. 6B**). Epithelial cells recovered from primary transplants were again expanded in FAD+Y+WS6 to obtain the cell numbers required for secondary transplantation (**Fig. 6A**). After 8 days, engraftment was detected in 5/5 second generation mice by IVIS imaging (**Fig. 6B**) and regeneration of the mouse trachea with ZsGreen-positive human epithelial cells was detected at day 49 (**Fig. 6C**). Tracheal regeneration was observed in histological assays of both first and second generation mice (**Fig. 6C-D**). Cells explanted from the mouse tracheas were enriched for ZsGreen+ cells and maintained epithelial cobblestone morphology (**Fig. 6E**). Colony formation assays before transplantation and following primary or secondary transplantation revealed maintenance of colony forming potential of ZsGreen-positive human basal cells (**Fig. 6F**).

**Figure 6:**
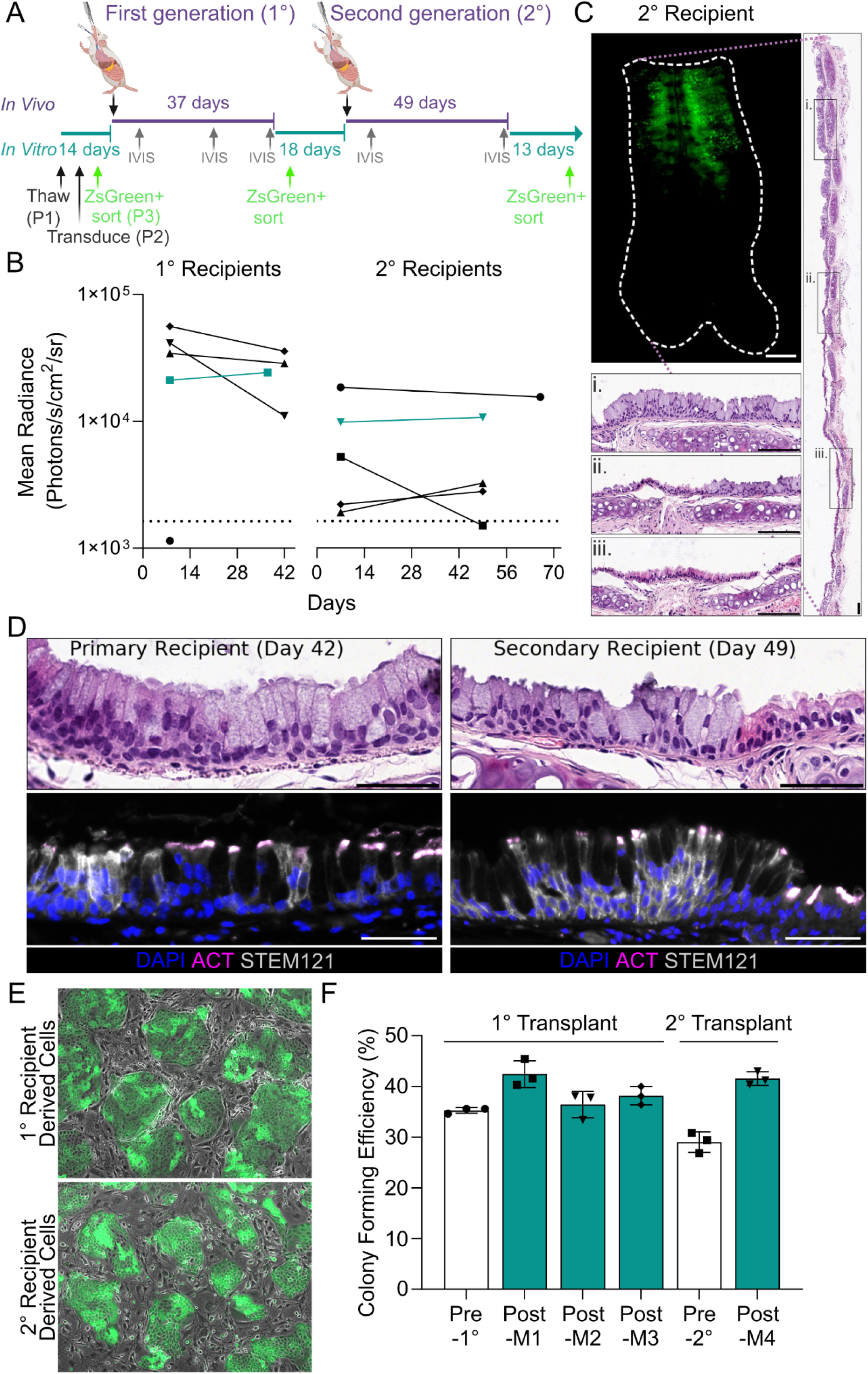
Serial tracheal transplantation of human basal stem cells expanded in FAD+Y+WS6. A) Schematic of serial transplantation of ZsGreen-luciferase transduced nasal epithelial basal cells in the trachea of immunodeficient NSG mice and monitoring by in vivo luciferase bioluminescence imaging (IVIS). B) Quantification of IVIS signal following cell engraftment (explanted animal tracheas are indicated in green) C) Whole-mount imaging of ZsGreen in a freshly excised trachea (scale bar = 1 mm) from a second generation mouse, 49 days following basal cell transplantation and hematoxylin and eosin staining i) majority human cell engraftment ii) mix of human cell engraftment and host mouse airway epithelium iii) host mouse airway epithelium (scale bars = 100 µm). D) Hematoxylin and eosin staining of sections from a mouse trachea after basal cell transplantation (upper) and immunofluorescence staining (acetylated tubulin, ACT, ciliated cells, magenta) and human origin (STEM121, grey). Scale bar = 50 µm. E) Phase-contrast and fluorescent imaging of flow-sorted ZsGreen-luciferase transduced nasal epithelial basal cells recovered from mouse tracheas. Scale bar = 100 µm. F) Colony formation assays (CFAs) to assess the effect of transplantation on clonogenicity of nasal basal epithelial cells. Cells transplanted into primary recipients were transduced at passage 2 and expanded for three passages prior to transplantation. Cells recovered from individual mice (M) were expanded for three passages prior to CFA. Pre-2° data is from cells recovered from M1 and expanded for five passages prior to CFA. Each point represents a single well replicate from one assay.

### Clonal genome editing of primary nasal airway basal cells for ciliary dyskinesia modeling

Many gene editing studies in epithelial cells rely on immortalized lines rather than primary cultures (Huang et al., 2022; Smits et al., 2023). While clonal basal cell cultures have enabled insights into genetic heterogeneity (Yoshida et al., 2020), their expansion and manipulation remain inefficient. In most cell culture systems, clonal cells are difficult to passage, cryopreserve, and differentiate. As a result, in cases where primary cells are used, most approaches involve bulk populations that contain a mixture of edited and unedited cells (Cattaneo et al., 2024; Chu et al., 2015; Niklander & Hunter, 2023; Rapiteanu et al., 2020).

Others use lentiviral systems with selectable markers, but these often result in persistent expression of Cas9 and guide RNAs (Fenini et al., 2018), limiting their suitability for therapeutic applications. To evaluate whether culture in FAD+Y+WS6 allows clonal cell expansion after gene editing, we used CRISPR–Cas9 genome editing to develop primary ciliary dyskinesia (PCD) disease models (**Fig. 7A**). First, we targeted *DNAH5*, the most commonly mutated gene in PCD (15-21% of cases (Damseh et al., 2017)), which disrupts outer dynein arm (ODA) function and produces a well-defined phenotype of ciliary immotility (Wallmeier et al., 2020). Nucleofection of CRISPR-Cas9-sgRNA ribonucleoprotein complexes introduced gene edits in bulk nasal basal cell cultures. Sanger sequencing showed a knockout efficiency of 93.1% and in parallel GFP plasmid nucleofections, an 89% efficiency was achieved (**Supplementary Fig. 13A**). As expected, edited bulk cultures displayed a mixture of beating and static cilia (**Supplementary Video 1**). Consistent with this, both normal and ODA-defective cilia were present within individual ALI cultures by transmission electron microscopy (TEM; **Supplementary Fig. 13B**).

Clones were isolated by single-cell sorting and transferred through 96-, 24-, and 6-well formats, with some attrition at each stage (**Fig. 7B**). Overall, 7% of of sorted single cells expanded in 6-well plates, and 14 of 19 (73.7%) of these were *DNAH5* knockouts (**Fig. 7C**). Clonal cultures retained the capacity to differentiate into both mucosecretory and ciliated cell lineages (**Fig. 7D**). Their epithelial integrity and barrier function, assessed by TEER, were comparable to non-PCD and PCD ALI cultures (**Supplementary Fig. 13C**). In clonal knockout ALI cultures, DNAH5 expression was undetectable by immunofluorescence (**Fig. 7E**). TEM confirmed ODA defects in all cilia assessed from clonal knockout cell cultures (100% ODA defects based on 50 cilia cross sections for *DNAH5* KO clones; **Fig. 7F**). Ciliary beat frequency measurements confirmed no detectable motility, faithfully recapitulating the PCD phenotype for *DNAH5* mutation (**Fig. 7G**). A second experiment targeting the *DNAI2* gene had similar results. Sanger sequencing showed knockout efficiency of 80.2% and 61.7% in cells nucleofected with sense and antisense single-stranded oligodeoxynucleotide (ssODN) donor template, respectively. Moreover, 66% nucleofection efficiency was achieved with a GFP plasmid (**Supplementary Fig. 14A**) and analysis of clonal cultures revealed that 22 of 37 (59.5%) clones were *DNAI2* knockouts (**Supplementary Fig. 14B**). TEER readings confirmed that their epithelial integrity and barrier function were comparable to non-PCD and PCD ALI cultures (**Supplementary Fig. 14C**). TEM again showed that only a subset of cilia were defective in bulk cultures (**Supplementary Fig. 14D**), while clones had either consistently intact or consistently defective ciliary ultrastructure (**Supplementary Fig. 14E**). Western blot confirmed loss of DNAI2 in knockout clonal cultures (**Supplementary Fig. 14F**). Following clonal expansion, neither wild-type or *DNAI2*-edited cells showed chromosomal abnormalities (**Supplementary Fig. 14G**).

**Figure 7:**
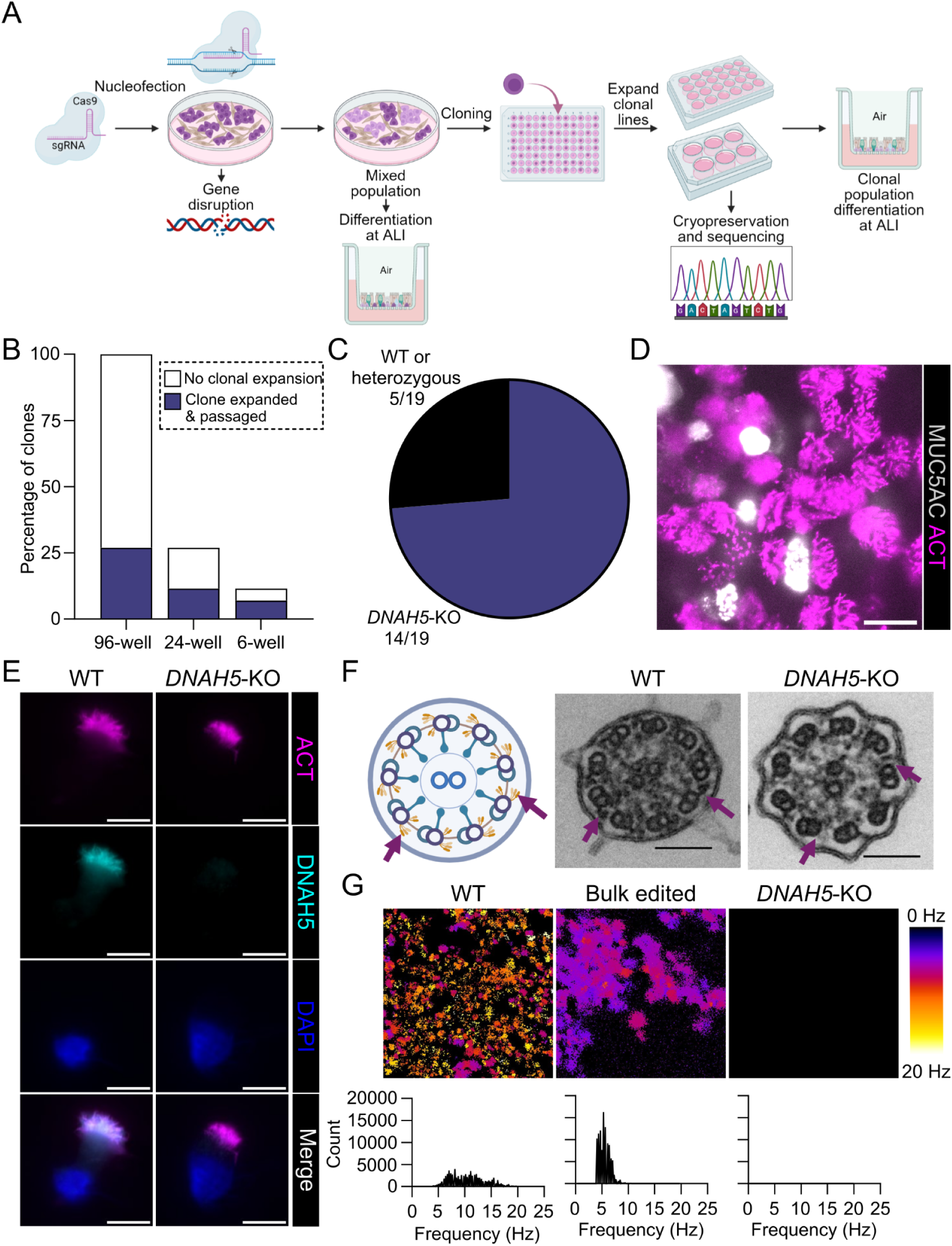
Primary ciliary dyskinesia disease modeling through clonal basal cell culture. A) Schematic representation of the workflow for epithelial basal cell expansion in FAD+Y-27632(Y)+WS6, CRISPR-Cas9 genome editing and differentiation in air-liquid interface (ALI) cultures. B) Quantification of the percentage of basal cell clones that reached 60% confluence from 285 wells and were passaged at each stage of expansion between 96-, 24- and 6-well plates (representative of three clonal culture experiments with this cell donor). C) Quantification of Sanger sequencing results showing the proportions of *DNAH5*-knockout (KO) or wild-type (WT)/heterozygous clones. D) Representative immunofluorescence staining of a basal cell clone (passage six, four passages after clonal cell sorting) after differentiation at ALI (acetylated tubulin, ACT, magenta; MUC5AC, grey). Scale bar = 20 µm. E) Representative immunofluorescence staining of individual ciliated cells from WT and *DNAH5*-KO clones differentiated at ALI (acetylated tubulin, ACT, magenta; DNAH5, cyan; DAPI, blue; n = 4 WT clones, n = 3 KO clones; passage seven, five passages after clonal cell sorting). Scale bar = 10 µm. F) Transmission electron microscopy of ciliary cross-section from WT and *DNAH5*-KO clones differentiated at ALI (passage seven, five passages after clonal cell sorting). Arrows show normal outer dynein arms in a WT clone and loss of ODA in *DNAH5*-KO clones (40,000× magnification, scale bar = 100 nm, n = 1 WT clone, n = 3 KO clones). G) Ciliary beat frequency analysis of differentiated ALI cultures derived from WT clones, the bulk population containing both *DNAH5*-KO and unedited cells, and *DNAH5*-KO basal cell clones (n = 3 WT and *DNAH5*-KO clones, and n = 3 experimental replicates of the bulk population). Coverage and beat frequency of moving cilia are displayed for representative fields of view (top) and frequency distributions between the 4-20 Hz analysis cut-offs (bottom).

## Discussion

Our data establish EpMED (FAD supplemented with Y-27632 and WS6, used in 3T3-J2 feeder-cell co-culture) as a practical approach for expanding primary human airway basal cells from low-input nasal brush biopsies. EpMED improved initial colony formation, cumulative expansion and culture longevity, maintained clonogenic capacity and supported mucosecretory and ciliated cell differentiation at later passages. Expanded cells retained the capacity to regenerate airway epithelium through serial transplantation, while clonal expansion following genome editing enabled primary cell-based primary ciliary dyskinesia modeling in isogenic wild-type and knockout cells.

Previous studies have shown that FAD-based 3T3-J2 feeder-cell co-culture can support long-term expansion of primary (pseudo)stratified epithelial cell types (Corradini et al., 2016; Droz-Georget Lathion et al., 2015; Pellegrini, Golisano, et al., 1999), including airway cells (Adamo et al., 2025). Here, we sought to improve expansion from minimally invasive nasal brush biopsies, which provide few initial basal cells. Previous studies have recovered approximately 1 × 10^6^ total cells per brushing, although yield varies with technique (Fawcett et al., 2023; Stokes et al., 2014). Adding WS6 to FAD+Y medium increased initial colony formation, accelerated expansion and prolonged culture duration, while maintaining colony formation and differentiation potential at later passages. These improvements facilitate experiments requiring large numbers of basal cells, such as parallel differentiations, or clonal genetic manipulation. EpMED also supported greater expansion and total yield than PneumaCult™-NGEx in a matched comparison. While the latter does not require feeder cells, WS6 did not eliminate the requirement for 3T3-J2 feeder cells in the FAD system; although cells could be passaged a number of times in the absence of feeder cells, their morphology and longevity were adversely affected.

Airway cells typically decline in their differentiation capacity and ion channel function over serial passage. However, nasal airway basal cells expanded in EpMED retained basal cell markers, differentiated into ciliated and mucosecretory lineages in ALI and organoid assays, and retained epithelial barrier, ciliary function and ion transport at passage 10. A higher degree of mucosecretory differentiation at passage 10 compared to passage 4 was observed in both FAD+Y and FAD+Y+WS6, which is consistent with an increasing propensity of basal cells to form mucosecretory cells over *ex vivo* culture observed in other culture media (Murthy et al., 2025). In an exceptional donor culture that was continuously passaged for over 18 months, differentiation to mucosecretory and ciliated cells was observed at passage 38, but ciliated differentiation was not observed at passage 100. Previous studies have identified declining ion channel function during the first five passages as a limitation of Y-27632-containing cultures (Schmidt et al., 2022), suggesting that EpMED culture could improve the useful lifespan of basal cells in this context without the use of low oxygen conditions (Peters-Hall et al., 2018).

The molecular mechanisms through which 3T3-J2 feeder cells, Y-27632 and WS6 enhance airway basal-cell expansion remain unresolved, but our data suggest that WS6 activity depends on cellular and culture contexts. Previous reports described its pro-proliferative effects on pancreatic alpha and beta cells (Boerner et al., 2015; Shen et al., 2013). In our experiments, the principal benefit of WS6 supplementation was in sustained 2D primary basal cell expansion. WS6 improved epithelial cell colony formation in nasal, tracheal, bronchial, small airway, and thymus primary cell cultures, with a similar trend observed for epidermal keratinocytes. Results in primary epithelial organoid cultures were mixed, with increased organoid size in small airway organoids, reduced size in alveolar organoids and no effect on the size of gastric organoids. *BMI1*-transduced nasal cells, but not those immortalized with hTERT/CDK4, increased proliferation in response to WS6. Two lung cancer cell lines did not. WS6 has been reported to act through targets including PA2G4 and IKKε (Shen et al., 2013), but the contributions of these proteins to airway basal stem cell phenotype are not yet known. We have not attempted to expand primary human tumor material in EpMED, but previous studies using Y-27632 alone have reported mixed results, with low efficiency and preferential expansion of normal epithelial stem/progenitor cells (Hynds, Ben Aissa, et al., 2018; Yu et al., 2017). Tumor-directed applications should therefore include enrichment for malignant cells and verification of expected genomic alterations early in and throughout culture.

Previous work demonstrated the capacity of early passage primary human airway basal cells or induced pluripotent stem cell-derived basal cells to engraft in polidocanol-conditioned NSG mouse trachea (Ma et al., 2023). One of the major challenges identified by Ma et al. and addressed in our study is the requirement for producing large numbers of primary human cells for translational applications. Our system can achieve this whilst preserving regenerative capacity over a greater number of cell passages than previously reported (Ma et al., 2023). Human cells engrafted in mice differentiated to mucosecretory and ciliated cell lineages, could be recovered and re-cultured, and demonstrated comparable clonogenic capacity in culture to before transplantation. Following a second *in vitro* expansion, cells were able to regenerate the airway epithelium of secondary recipient mice.

Previous studies have investigated airway epithelial gene function and disease mechanisms using CRISPR-Cas9 gene knockout in primary cells, including for the cell adhesion molecule *MUC18* (Chu et al., 2015), the mucosecretory cell regulator *SPDEF* (Koh et al., 2020), the regulators of ciliogenesis *GRHL2* (Gao et al., 2015) and *FOXJ1* (Rapiteanu et al., 2020; Zaragosi et al., 2024) and the rhinovirus C receptor *CDHR3* (Everman et al., 2019). Modified conditional reprogramming conditions allowed clonal expansion of *CFTR*-edited human bronchial epithelial cells with retained differentiation capacity for ion channel investigations (Peters-Hall et al., 2018). In our study, while only a small proportion of cells were capable of clonal expansion, the ability to subculture from individual cells allowed us to similarly separate unedited wild-type cells from *DNAI2*-or *DNAH5*-knockout cells with distinct editing genotypes. Clonal expansion resolved the mixed ultrastructural and motility phenotypes that were seen in bulk edited populations and allowed direct comparison of isogenic clonal cultures with defined genotypes. In the future, the ability to derive clonal cultures may also offer safety advantages for epithelial cell therapy by enabling preclinical characterization of viral integration sites prior to transplantation (Droz-Georget Lathion et al., 2015).

Improved expansion conditions for airway basal cells may ultimately support autologous epithelial cell therapy: primary epithelial cultures are already used clinically for severe burns and limbal stem-cell deficiency (Pellegrini, Ranno, et al., 1999; Rama et al., 2010), and *ex vivo* gene-corrected epidermal cells have been used successfully in epidermolysis bullosa (Kueckelhaus et al., 2021). Based on the expansion rates measured here, we estimate that a nasal brushing could yield the number proposed for whole-organ tracheal bioengineering in 12-18 days (approximately 70 million basal cells (Butler et al., 2016)). However, EpMED depends on fetal bovine serum and 3T3-J2 mouse feeder cells, which complicates the transition to a GMP-compatible production process. Moreover, the present study used mitomycin C-treated feeder cells, whereas lethally irradiated feeder cells would be required for a GMP process because residual mitomycin C and/or inadequate feeder inactivation could create a patient safety risk (Pellegrini et al., 2018).

A number of feeder-free systems have emerged as alternatives to 3T3-J2 co-culture but these have limitations for translational applications. While the NGEx reference medium used here is ‘defined’, its composition is not known to the user, limiting mechanistic interpretation and additional modifications. Dual SMAD inhibition can improve basal cell expansion (Mou et al., 2016), although this approach relies on coating culture vessels with conditioned medium from a rat bladder cancer cell line. Other laboratories have developed in-house custom media that support serum-free and feeder-free expansion of airway basal cells (Dreyer et al., 2026; Jiang et al., 2026) but characterization of long-term genome stability and long-term *in vivo* regeneration potential are limited for these methods.

As discussed above, further work is required to establish the mechanisms by which 3T3-J2 feeder cells, Y-27632 and WS6 combine to enable extensive epithelial cell expansion. While our long-term expansion, functional and *in vivo* work were performed in nasal basal cells, colony formation experiments suggest that the method may be relevant to other (pseudo)stratified epithelial cells. There also remains substantial heterogeneity in the expansion potential of individual donor samples. The origin of this heterogeneity remains unresolved but could relate to donor characteristics, such as donor age (Farr et al., 2026), and/or the number of basal cells in the initial biopsy with high *ex vivo* clonogenic potential. While low-pass whole-genome sequencing and tumorigenicity experiments herein are reassuring, they do not establish comprehensive genomic stability or clinical safety; deeper genomic characterization and longer safety studies are required.

Our work establishes a practical, robust *ex vivo* culture system for expansion of primary airway basal cells from limited initial clinical material for *in vitro* and *in vivo* experimentation. Cells retain progenitor cell potential, generate functional differentiated progeny, and can be clonally expanded and differentiated in ALI cultures following genome editing.

## Methods

### Cell culture

#### 3T3-J2 fibroblast culture and feeder layer preparation

3T3-J2 mouse embryonic fibroblasts (Kerafast) were maintained in DMEM with sodium pyruvate (Gibco, 41966) supplemented with 7% bovine serum (Hyclone, SH30072.04; lot AE29427271), 100 U/mL penicillin, and 100 μg/mL streptomycin (Sigma-Aldrich, P0781-100ML). Cultures were maintained at 37 °C in a humidified 7.5% CO_2_ incubator, with medium replaced three times per week.

To prepare feeder layers, cells were mitotically inactivated by treatment with 4 μg/mL mitomycin C (Sigma-Aldrich, M4287, 0.4 mg/mL stock in PBS) for 3 h. After washing and trypsinization, cells were seeded at 20,000-30,000 cells/cm² in fresh 3T3-J2 cell culture medium. Epithelial cells were added the following day.

#### Primary epithelial 2D cell culture

Human basal epithelial cells were obtained from nasal, tracheal, bronchial, lung, thymus and skin tissues with patient consent (Table S1). Ethical approval was obtained through National Research Ethics Committees (Living Airways Biobank, REC 24/NW/0168, nasal cells; REC 18/SC/0514, tracheal, bronchial, small airway cells; REC 11/LO/1522, skin and gastric cells; REC 07/Q0508/43, thymus cells).

Tissues were transported to the laboratory in transport medium consisting of αMEM (Gibco, 22571020) supplemented with 100 U/mL penicillin, and 100 μg/mL streptomycin (Merck, P0781), 10 μg/mL gentamicin (Gibco, 15710-049) and 250 ng/mL amphotericin B (Fisher, 10346503). Nasal brushings were performed using cytology brushes (Medical Packaging Corporation, CYB-1). To generate single cell suspensions, the brushes were incubated in 0.1% trypsin/EDTA (Sigma-Aldrich, 59418C) in RPMI (Gibco, 21875) for 20 min at 37°C with vigorous shaking every 2 min. The cell suspensions were transferred to new 15 mL tubes and the brushes washed with RPMI medium. After this, the trypsin was neutralized with fetal bovine serum (FBS; Gibco, A5256701; lot 2575638). The cell suspensions were centrifuged at 300 × g for 5 min and resuspended in red blood cell lysis buffer (Life Technologies, 00-4333-57) for 10 min at room temperature. The cells were centrifuged again, washed with RPMI, centrifuged and resuspended in the relevant medium for cell seeding. Lung tissue was minced using a sterile scalpel for 5 min, this tissue and tracheal and bronchial biopsies were incubated in 16 U/mL dispase (Corning, 354235) in RPMI for 20 min at room temperature and then mechanically disrupted using sterile forceps. This was followed by incubation in 0.1% trypsin/EDTA for 30 min at 37°C and neutralization with FBS. The cells were filtered through a 100 µm strainer (Falcon, 352360) and single cell suspensions were centrifuged at 300 × g for 5 min and resuspended in the relevant medium for cell seeding.

FAD medium was prepared by combining 342 mL DMEM without pyruvate (Gibco, 41965062; 158 mL removed from a 500 mL bottle), 33.3 mL FBS and 112.5 mL Ham’s F-12 (Gibco, 21765029). The medium was then supplemented with the following: 5 mL penicillin-streptomycin (10,000 U/mL penicillin and 10 mg/mL streptomycin stock in 0.9% NaCl, 100 U/mL and 0.1 mg/mL final), 5 mL adenine (Sigma-Aldrich, A3159-5G; 17.9 mM stock in 0.1 M HCl, 179 μM final), 0.5 mL gentamicin (Life Technologies, 15710049; 10 mg/mL stock, 10 μg/mL final), 40 μL hydrocortisone (Sigma-Aldrich, H0888-1G; 6.25 mg/mL in ethanol, 0.5 μg/mL final), 495 μL triiodothyronine (Sigma-Aldrich, T6397-100MG; 2.02 μM stock in PBS, 2 nM final), 0.5 mL insulin (Merck, I0516; 5 mg/mL in ∼12.5 mM HEPES, 5 μg/mL final), 4.2 μL cholera toxin (Sigma-Aldrich, C8052-1MG; 11.9 μM stock in water, 0.1 nM final), and 50 μL EGF (PeproTech, AF-100-15-500ug; 100 μg/mL stock in 0.1% BSA/PBS, 10 ng/mL final). Where indicated, 5 μM Y-27632 (Tebu Bio, T1725; 500 μL of 5 mM stock in distilled water) and/or 100 nM WS6 (APExBIO, B4880; 50 μL of 1 mM stock in DMSO) were also added. This epithelial cell medium (‘EpMED’; FAD containing 5 μM Y-27632 and 100 nM WS6) is now our routine cell culture medium for establishing and maintaining basal epithelial cell cultures. We prepare 10x concentrated stocks, FBS, penicillin-streptomycin and the supplements listed above as a 10× EpMED stock, stored at −80°C in 10 mL or 50 mL aliquots and thawed into 90 mL or 450 mL DMEM without pyruvate:Ham’s F-12 (3:1) as required, respectively. This gives equivalent final concentrations (Table S2).

PneumaCult™-NGEx Medium (STEMCELL Technologies, 05001) was prepared according to the manufacturer’s instructions (Basal Medium + 50X Supplement + Hydrocortisone Stock Solution (STEMCELL Technologies, 07925)) and supplemented with 100 U/mL penicillin, 100 μg/mL streptomycin and 10 μg/mL gentamicin. Tissue culture plastic was coated with 1 % collagen I (STEMCELL Technologies, 07001) in PBS for 30 min at room temperature and left to air dry for 5 min before cell addition.

Epithelial cells were cultured at 37**°**C and 5% CO_2_ with three medium changes per week. Epithelial cells can be enriched from co-cultures using differential trypsinization; 3T3-J2 feeder cells have a greater sensitivity to trypsin than primary basal cells allowing a first trypsin-wash cycle to remove only the 3T3-J2 cells, and a subsequent trypsin step to remove the more firmly attached epithelial cells. Epithelial cells were counted following differential trypsinization and population doublings (PD) were calculated as PD = 3.32 × [log(cells harvested/cells seeded)].

#### Primary small airway and alveolar organoid culture

Human small airway (Sachs et al., 2019) and alveolar organoid (Dost et al., 2026) cultures were derived from distal lung tissue (Diakonessen Hospital Utrecht, The Netherlands) as previously described. Cells were plated as single cell suspensions at 500 cells/µL (small airway) or 300 cells/µL (alveolar) in 10 µL 80% Basement Membrane Extract (R&D Systems, 3533-005-02) at the bottom of a 96-well organoid culture plate (STEMCELL Technologies, 200-0562).

#### Primary gastric organoid culture

Endoscopic mucosal biopsies taken from the fundus, body and antrum of the stomach were dissociated to single cells in TrypLE Express (Thermo Fisher Scientific, 12605010) for 5 min and cultured in 12-well plates (Thermo Fisher Scientific, 10098870) in 30 µL droplets of Matrigel (Corning, 354230). Gastric organoid medium consisted of Advanced DMEM F-12 (Thermo Fisher Scientific, 12634) supplemented with 10 mM HEPES (Thermo Fisher Scientific, 15630080), 2 mM Glutamax (Thermo Fisher Scientific, 35050061), 1% penicillin-streptomycin, 100 µg/mL primocin (Thermo Fisher Scientific, NC9392943), 1× B-27 supplement minus vitamin A (Thermo Fisher Scientific, 12587010), 1.25 mM n-acetylcysteine (Sigma-Aldrich, A9165), 100 ng/mL Wnt-3A (Peprotech, 315-20), 500 ng/mL R-spondin 1 (Peprotech, 120-38), 100 ng/mL noggin (Peprotech, 120-10C), 50 ng/mL human EGF (Thermo Fisher Scientific, PMG8043), 10 nM gastrin (Sigma-Aldrich, G9020), 3 µM CHIR99021 (Tocris, 4423), 5 µM A83-01 (Sigma-Aldrich, SML0788) and 200 ng/mL FGF10 (Peprotech, 100-26). This medium was supplemented with 10 µg/mL Y-27632 (Tocris, 1254) for the first 24 h. Organoids were passaged by TrypLE and manual disaggregation with a P1000 pipette and split 1:3-15.

#### Primary hematopoietic stem and progenitor cell culture

Human cord blood CD34+ cells (Anthony Nolan) were cultured in SFEM II (STEMCELL Technologies, 09655) supplemented with 150 ng/mL human stem cell factor (Peprotech, 300-07-10UG), 150 ng/mL human FLT3-ligand (Peprotech, 300-19-10UG), 20 ng/mL human thrombopoietin (Peprotech, 300-18-10UG) and 1% penicillin-streptomycin. Where indicated, the medium was supplemented with 10 nM, 100 nM or 1 μM WS6. Cells were cultured at 37°C and 5% CO_2_ for 7 d with a medium change on day four.

#### Cell line culture

HBEC3-KT cultures were maintained in Keratinocyte-SFM (Gibco, 17005042); A549 and H520 cells were maintained in RPMI 1640 (Gibco, 21875034) supplemented with 10% FBS, 1 mM sodium pyruvate (Gibco, 11360070), 100 U/mL penicillin, and 100 μg/mL streptomycin. *BMI1*-transduced nasal cells (Maeshima et al., 2024) were maintained in PneumaCult™-Ex Medium (STEMCELL Technologies, 05008) on tissue culture plastic coated with 1% PureCol (Advanced BioMatrix, 5005) in PBS for 30 min at room temperature and left to air dry for 5 min. All cultures were maintained at 37°C and 5% CO_2_, with medium changes three times per week.

#### Nasal organoid differentiation assay

To generate 3D nasal airway organoids, basal cells were enzymatically dissociated and counted. Organoid medium consisted of 50% BEBM (Lonza, 3171) and 50% DMEM (Gibco, 41966) supplemented with BEGM additives (Lonza, 4175; excluding triiodothyronine, gentamicin, amphotericin B, and retinoic acid). Retinoic acid (100 nM; Sigma-Aldrich, R2625, stock 10 mM in ethanol) was freshly added before use. 5 μM Y-27632 was added to the organoid medium for cell seeding. Ultra-low attachment 96-well plates (Corning) were pre-coated with 30 μL of 25% Matrigel (Corning, 354230; growth factor–reduced) in organoid medium and incubated at 37°C for 30 min. Then, 2,500 basal cells were seeded per well in 65 μL of 5% Matrigel. Cultures were fed by adding 70 μL of fresh medium on days 3, 10, and 17. At 21 d, whole-well images were taken at 10× magnification with a Nikon Eclipse Ti2-E inverted microscope and tracheosphere size was quantified using OrgaQuant(Kassis et al., 2019). R was used to remove aberrant detection, including objects within 110 pixels of the image edges. Predicted organoid volume was calculated as (*V* = (4/3) * π * *r*³) and the mean was taken from all organoids detected per condition. Organoids were harvested with ice-cold PBS, and centrifuged at 300 × g for 5 min. Organoids were fixed in 4% PFA for 30 min on ice, washed in PBS, and embedded in HistoGel^TM^ (Epredia, HG-4000-012) for paraffin processing.

#### CellTiter-Glo assays

For primary 2D cell cultures, compounds and DMSO were added to the black-walled 384-well assay plates (Thermo Fisher Scientific, 142761) using an ECHO 550 liquid handler (Labcyte) or in 5 µL FAD medium. 2,000 cells/well were then added in FAD medium using a microplate washer/dispenser (BioTek EL406, Agilent) or using a multichannel pipette under sterile conditions. After 4 or 6 d of culture, the medium was removed and the cell number was determined using the CellTiter-Glo Luminescent Cell Viability protocol (Promega). After 20 min, bioluminescence was measured using a Envision II plate reader (PerkinElmer) or a VANTAstar plate reader (BMG LABTECH, 1 s integration time).

For cell lines, 1,000 cells/well were seeded in black-walled 96-well plates (Corning, 3603), after 48 h compounds were added in cell culture medium (0.1% DMSO), after 48 h the medium was removed and the CellTiter-Glo Luminescent Cell Viability protocol was performed as above.

For small airway and alveolar organoids, 100 µL organoid medium was supplemented with 10 nM, 100 nM, 1 µM WS6 or DMSO as a vehicle control (small airway; 1%, alveolar; 0.1%) and the medium was refreshed every 2-4 d. Images were taken at 14 d and the CellTiter-Glo 3D Cell Viability Assay (Promega, G9681) was performed according to the manufacturer’s protocol. 100 µL CellTiter-Glo reagent was added to the wells and the plate was put on a Thermo Fisher Scientific Multidrop Combi+ plate shaker for 15 min at room temperature. After an additional incubation time of 15 min at room temperature, 100 µL of the solution was transferred to a black bottom/black wall plate (Greiner Bio-One, 655076) and luminescence was measured (Tecan Spark, 500 ms integration time). Measurements from each well were normalized to the mean of measurements of three replicate wells taken in the same way on day 0, immediately after plating the organoid lines.

For gastric organoids, 500 µL organoid medium was supplemented with 10 nM, 100 nM, 1 µM WS6 or 0.1% DMSO as a vehicle control and the medium was refreshed every 3 d. Images were taken at 7 d, the organoid medium was removed and replaced with 30 µL ice-cold RPMI. 100 µL CellTiter-Glo reagent was added and the plate was shaken at 60 rpm on a see-saw shaker (Stuart) for 15 min. 100 µL of the solution was transferred to a black wall plate (Corning, 3603) and luminescence was measured on a VANTAstar plate reader (1 s integration time). Measurements from each well were normalized to the mean of measurements of three replicate wells taken in the same way on day 0, immediately after plating the organoid lines.

#### Colony formation assays

To assess colony forming efficiency (CFE), epithelial basal cells were seeded at 1,000 cells per well with technical triplicates into six-well plates that had been seeded with 3T3-J2 feeder cells the day prior. After 7 d, plates were fixed in 4% paraformaldehyde (PFA) for 10 min at room temperature and stained with 1% crystal violet solution (Sigma-Aldrich) for a further 10 min. Plates were washed in water and air-dried overnight. Colonies were counted manually under a brightfield microscope, classifying all coherent colonies of more than 10 cells as a colony. CFE was calculated as: (number of colonies / number of seeded cells) x 100. Plates were imaged using an Epson Perfection V600 flatbed scanner.

#### Histology processing

Chamber slides were fixed in 4% PFA for 10 min, washed with PBS and stored in PBS prior to staining. Fixed, histogel-embedded organoids and ALI cell cultures, and formalin-fixed tissues were processed in a Leica TP1050 tissue processor. Samples were embedded in type 6 paraffin wax (Epredia, 8336) using an embedding station (Sakura Tissue-TEK TEC) and 5 µm sections were cut on a Microm HM 325 microtome and stored at 4°C before use.

Hematoxylin and eosin (H&E) staining was performed on sections using an automated staining system (Sakura TissueTek DRS) and imaged on a Nanozoomer Digital Pathology (Hamamatsu). Histology images were collated and presented from whole-slide image files using PATHOverview (Pearce et al., 2023) (available on GitHub https://github.com/EpiCENTR-Lab/PATHOverview).

#### Immunofluorescence staining

Slides were dewaxed using an automated protocol, washed in PBS, and a hydrophobic ring was drawn around the sample using an ImmEdge pen (Vector Laboratories, H-4000). Sections were blocked with 1% bovine serum albumin (BSA; Merck, 1.12018.0100), 5% normal goat serum (NGS; Gibco, 16120064) and 0.1% Triton X-100 (Sigma-Aldrich, X-100) in PBS for 1 h at room temperature. Primary antibodies (Table S3) were diluted in blocking buffer and applied to slides overnight at 4°C. Slides were washed twice in PBS. Species-appropriate secondary antibodies conjugated to Alexa Fluor dyes (Table S4) were diluted 1:1,000 in 5% NGS, 0.1% Triton X-100 in PBS and applied to slides for 3 h at room temperature in the dark. 200 ng/mL DAPI (Sigma-Aldrich, D9542) in PBS was applied to the slides for 20 min. Slides were washed twice in PBS and a coverslip was applied manually with Immu-Mount (Thermo Fisher Scientific, 9990402). Whole-mount ALI cultures were transferred to 1.5 mL Eppendorf tubes and blocked in PBS containing 2% BSA, 5% NGS, and 0.1% Triton X-100 for 1 h at room temperature at 20 rpm on a see-saw shaker. Primary antibodies (Table S3) were diluted in blocking buffer and incubated overnight at 4°C at 60 rpm on a mini orbital shaker (Stuart, SSM1). After three 5-min PBS washes on the see-saw shaker, secondary antibodies (Table S4) were diluted 1:1,000 in 2% BSA, 5% NGS, 0.1% Triton X-100 in PBS and applied for 3 h at room temperature in the dark on the see-saw shaker. DAPI (Sigma-Aldrich, D9542) was applied at 100 ng/mL for 20 min in the dark on the see-saw shaker. Inserts were washed twice in PBS and once in distilled water, then mounted with Immu-Mount within imaging spacers (Grace Bio-Labs, 654006).

For staining of individual ciliated cells, ALI cultured cells were scraped off transwell inserts, collected in PneumaCult™-ALI medium (STEMCELL Technologies, 05001) and dissociated by pipetting. Cells were left to air dry on slides (Epredia, 5991056) and stored at 4°C. All subsequent incubation steps were performed in the dark at room temperature. Cells were fixed in 4% PFA in PBS for 15 min and washed twice with 0.1% Triton X-100 in PBS (wash buffer used for all washes). Slides were blocked in PBS with 5% w/v milk powder (Marvel) and 0.1% Triton X-100 for 1 h, washed twice and incubated with anti-DNAH5 primary antibody (Table S3), diluted 1:200 in wash buffer, for 2 h. After two washes, slides were incubated with anti-acetylated tubulin primary antibody (Table S3), diluted 1:2500 in wash buffer, for 45 min. Slides were washed twice and incubated in secondary antibodies (Table S4), diluted in wash buffer, for 30 min. After antibody incubations, slides underwent three 1-min washes. Slides were mounted using ProLong Gold antifade reagent with DAPI (Invitrogen, P36935) and left to cure overnight before imaging on a Nikon Eclipse TE2000-U inverted microscope.

All other immunofluorescence images were acquired using a Nikon Eclipse Ti2-E inverted microscope.

#### Air-liquid interface cultures

Semi-permeable membrane supports (Pore size 0.4 µm; Corning, 3470 or Sarstedt Ltd, 83.3932.041) were coated with 50 µg/mL rat tail collagen I (Corning, 354236) in 0.02N acetic acid (Sigma-Aldrich, 1.00063.1011). After 1 h at room temperature, the liquid was removed and the membranes washed twice with PBS. They were left to air dry for 15 min. To establish ALI cultures, airway basal cells were seeded onto the collagen-coated semi-permeable membrane supports (2 x 10^5^ cells in 200 µL epithelial growth medium, Fig. 7 and Supplementary Fig. 13 and 14 3 x 10^5^ cells in 250 µL epithelial growth medium, Fig. 4 and Supplementary Fig. 10 and 12). 500 µL epithelial growth medium was added to the basolateral side. After 24 h (at confluence), medium was removed from the apical side and replaced basolaterally with PneumaCult™-ALI medium (STEMCELL Technologies, 05001) supplemented with 4 µg/mL heparin (Sigma-Aldrich, H4784) and 0.48 µg/mL hydrocortisone (Sigma-Aldrich,H0888). Medium was refreshed three times per week. Mucus was removed by gentle apical washing with PBS once a week. ALI cultures were maintained for 28-35 d before analyses. Transepithelial electrical resistance (TEER) was measured using an EVOM manual resistance meter (World Precision Instruments). Mucus was removed by gentle apical washing with PBS and measurements taken with 200 ul of PBS (pH 7.2, Ca- Mg-, Gibco, 20012-019) added to the apical compartment. Three readings were taken from each insert once resistance had stabilized (5-10 s) and triplicate inserts were measured for each condition.

#### Transepithelial ion transport

To establish ALI cultures for transepithelial ion transport analysis, 1 × 10^6^ airway basal cells were seeded onto collagen-coated Snapwell membranes (Corning, 3407) in 300 µL epithelial growth medium. ALI cell culture was carried out as above with 1.5 mL medium in the basolateral side. At 30 d, transepithelial ion transport was measured using the Ussing chamber technique with symmetrical bicarbonate buffered salt solution (in mM): NaCl, 117; NaHCO_3_, 25; KCl, 4.7; MgSO_4_, 1.2; KH_2_PO_4_, 1.2; CaCl_2_, 2.5; D-Glucose, 11 (equilibrated with 5% CO_2_ to pH 7.3-7.4). The solution was maintained at 37°C, bubbled with 21% O_2_/5% CO_2_ gas mix. The following drugs were applied: amiloride (apical, 100 μM) to inhibit the epithelial sodium channel (ENaC); forskolin (bilaterally, 10 μM) to stimulate CFTR; CFTRinh-172 (apical, 10 μM) to inhibit CFTR; ATP (apical, 100 μM) to stimulate calcium-activated chloride channels and ouabain (basolateral, 1 mM) as previously described(Rose et al., 2025; Woodall et al., 2023; Woollhead et al., 2005). All drugs/chemicals were obtained from Sigma-Aldrich. Data were collected and analyzed using LabChart 7 software (AD Instruments).

#### Functional analysis of cilia

For top-down ciliary beat frequency (CBF) analysis, ALI cultures were left to stabilize at 37°C for 30 min after PBS washing/TEER measurements. Fast time-lapse videos were acquired at 37°C with 5% CO₂, using a Nikon Eclipse Ti-E inverted microscope (Nikon, Japan) equipped with a Prime BSI Express camera and a Nikon Super Plan Fluor ELWD 20XC PH objective. For CBF analysis, we captured 512 frames at a frame rate of 86.95 frames per s. Cilia beat frequency and active area were quantified using the CiliaFreqMap ImageJ macro (Moulding, 2025) with the range 4-20 Hz.

To assess ciliary motility, differentiated epithelium was scraped from transwells using 200 µL of HEPES-buffered Medium 199 (Gibco, 22340-020), gently dissociated by pipetting, and transferred onto a glass slide for side-on visualization of the epithelium using the above Nikon Eclipse Ti-E inverted microscope. The remaining cell suspension was transferred onto a glass slide within a hydrophobic ring and allowed to air dry at room temperature. The slides were then fixed in 4% PFA for 15 min and immunofluorescence staining was performed as described for single ciliated cells, using 0.1% Triton X-100 (Sigma, X100-500ML) in PBS as wash buffer and 5% milk powder (Marvel) in 0.1% Triton X-100 PBS as blocking buffer. Slides were mounted with ProLong Gold antifade reagent with DAPI (Thermo Fisher Scientific, P36935).

#### Transmission electron microscopy

For transmission electron microscopy (TEM) of ciliary axonemes, the ALI cultures were scraped from transwells using 200 µL of HEPES-buffered Medium 199 (Gibco, 22340-020) and fixed with 2.5% glutaraldehyde and 2% paraformaldehyde in 0.1 M Sorensen’s buffer (pH 7.3) for 24 h. The cells were postfixed in 1% aqueous solution of osmium tetroxide.

Following several washes, the tissue was dehydrated through a graded ethanol series and then propylene oxide (Merck). The epithelium was embedded in TAAB812 resin (TAAB Laboratory Equipment) and sections, 70 nm in thickness, were cut using a Leica UC7 Ultramicrotome (Leica Microsystems). These sections were then mounted on copper mesh grids and contrasted for 2 min with 4% uranyl acetate solution in methanol (VWR), followed by 2 min in lead citrate (Reynolds’ solution). Imaging was performed using a JEOL JEM-1400 transmission electron microscope (JEOL) operated at 120 kV, and digital images were acquired using a Xarosa camera and Radius software (EMSIS).

TEM quantification of outer dynein arm (ODA) defects was performed in accordance with established clinical diagnostic guidelines for primary ciliary dyskinesia (Shoemark et al., 2020). In brief, 50 individual ciliary axonemal cross-sections were assessed from different regions throughout each sample. ODA defects were defined as truncated, structurally abnormal, or absent outer dynein arms. Quantification was performed using the Cell Counter plugin in FIJI (O’Brien et al., 2016).

#### Western Blotting

Differentiated epithelium from ALI cultures was scraped from transwells in 200 µL PBS and centrifuged at 400 × g for 5 min. Cell pellets were resuspended in RIPA lysis buffer (Thermo Fisher Scientific, 89901) containing protease and phosphatase inhibitors (Thermo Fisher Scientific, 10025743) on ice. Lysates were stored at -20°C and thawed on ice before use. Protein concentration was quantified using a Pierce BCA Protein Assay kit (Thermo Fisher Scientific, 23225), following the manufacturer’s microplate protocol. The assay was performed with triplicate technical replicates, using a 1:10 dilution of protein samples.

Protein samples were denatured in Laemmli buffer (Sigma-Aldrich, S3401-1VL) with a 10 min incubation at 95°C. Denatured samples were loaded into gels and electrophoresed at 120 V for 75 min (DNAI2 Supplementary Fig. 14F: 4-12% Bis-Tris gels (Invitrogen, NW04125BOX)) and MOPS SDS running buffer (Invitrogen, B0001). Protein was dry-transferred onto a nitrocellulose membrane (Invitrogen, IB33002) with an Invitrogen iBlot3 machine following manufacturer recommended transfer protocol based on target protein size.

Blots were blocked for 1 hr at room temperature in 5% milk powder (VWR Chemicals, 84615.0500) in tris-buffered saline (TBS; Cell Signaling Technology, 12498S) with 0.1% Tween 20 in TBS (TBST; Sigma-Aldrich, P9416-100ML), before overnight incubation with primary antibody (Table S5) at 4°C with rotation. Blots were washed in tris-buffered saline with TBST and transferred to HRP-conjugated secondary antibody (Table S5) for 2 h at room temperature. Blots were imaged after a 2 min incubation in Crescendo HRP substrate (Millipore, WBLUR0100) and chemiluminescence imaging on a Bio-Rad ChemiDoc MP imaging system.

#### Fluorescence-activated cell sorting for clonal cell culture

Single cell suspensions from digested bronchial biopsies and nasal brushings were stained in sorting buffer (1× PBS, 1% FBS, 25 mM HEPES (Gibco, 15630-049) and 1 mM EDTA (Invitrogen, AM9260G) with anti-CD45-PE (1:20 dilution; BD Pharmingen 555483), anti-CD31-PE (1:20 dilution, BD Pharmingen 555446), anti-EPCAM-APC (1:50 dilution, BioLegend 324208) antibodies for 15 min at 4°C. Cells were washed and resuspended in sorting buffer containing 1 μg/mL DAPI.

Single DAPI^-^CD45^-^CD31^-^EPCAM^+^ cells were sorted (BD FACS Aria Fusion) into wells of a 96-well plate, precoated with 50 µg/mL rat tail collagen I and mitotically inactivated 3T3-J2 feeder cells with 50 μL epithelial conditioned medium. Epithelial cell-conditioned medium was a 1:1 mix of fresh epithelial medium and sterile filtered medium (0.22 µm filter; SLS, B2B06412) that had been collected from proliferating epithelial cells 24 h after fresh medium was added (50-90% confluency), and supplemented with 10 ng/mL EGF, 5 μM Y-27632 and 100 nM WS6 where indicated. An additional 50 μL epithelial cell-conditioned medium was added 3 d post-sort. After 7 d and 10 d, medium was replaced with fresh epithelial medium. At 14 d, whole-well images were taken on a Leica Dmi8 microscope and the number of wells with a single colony were quantified.

#### Flow cytometry

Fully differentiated ALI cultures were harvested for flow cytometry to quantify ciliated and mucosecretory cell populations. Transwells were washed twice with pre-warmed PBS for 5 min at 37°C, excised, and transferred to 1.5 mL tubes containing 1 mL TrypLE Express dissociation buffer (Thermo Fisher Scientific, 12604013). Tissues were incubated at 37°C for 8 min, briefly vortexed, and returned to the incubator. This cycle was repeated once. Dissociated cells were transferred to DMEM supplemented with 10% FBS to neutralize enzymatic activity, centrifuged at 400 × g for 5 min at 4°C, and resuspended in PBS for counting and downstream processing.

A total of 1-2 x 10^5^ cells were pooled from all samples and divided into unstained, live/dead stained, and heat-shocked (65°C, 10 min) live/dead stained control conditions.

Cells were stained with LIVE/DEAD Fixable Dye (Thermo Fisher Scientific, L34992; 1:1,000 in PBS) for 20 min at 4°C, washed, and resuspended in FACS buffer (PBS, 1% BSA, 1 mM EDTA (Invitrogen, 15575-038). A third of these cells were distributed between fluorescence minus one (FMO) control wells. Following centrifugation at 400 × g for 5 min at 4°C, cells were stained with antibodies targeting cell surface antigens (Table S6) in 50 μL FACS buffer for 30 min at 4°C in the dark. After washing, cells were fixed in 100 μL 4% PFA for 20 min at 4°C, permeabilized in 150 μL permeabilization buffer (1:10 dilution in PBS; eBioscience, 00-8333-56), and stained with 50 μL intracellular antibodies in permeabilization buffer for 30 min at 4°C. Cells were washed twice in FACS buffer, resuspended in 100 μL FACS buffer, and analyzed on a CytoFLEX flow cytometer (Beckman Coulter). Compensation was performed using UltraComp plus eBeads (Invitrogen, 01-3333-41) and live/dead-stained cells.

#### CRISPR-Cas9 gene editing and clonal epithelial cell culture

Primary nasal basal cells were isolated from fresh nasal brushings or thawed from frozen stocks and expanded in WS6-containing medium on mitotically-inactivated 3T3-J2 fibroblasts.

For nucleofection, epithelial cells were harvested at ∼80% confluency by differential trypsinization to remove feeder cells. Cells were counted and resuspended at 2 x 10⁵ per reaction. Ribonucleoprotein (RNP) complexes were assembled by incubating 0.5 µL (30.5 pmol) Cas9 (IDT) and 0.5 µL (50 pmol) sgRNA (Table S7) at room temperature for 10 min. For *DNAH5* knockout, 1 µL of Cas9 electroporation enhancer (IDT) was added to improve nucleofection efficiency. For *DNAI2*, 1.5 µL of 100 µM donor template oligonucleotide introducing a premature stop codon was added; this ssODN served as both the HDR template and as an electroporation enhancer, yielding an HDR-mediated knockout. Both a sense and anti-sense ssODN were tested in separate nucleofections to maximise editing efficiency.

Nucleofection was performed using the 4D-Nucleofector System (Lonza) with the P3 Primary Cell kit. Each reaction included 2 x 10⁵ cells resuspended in 20 µL of P3 buffer (16.25 µL nucleofector solution + 3.75 µL supplement 1), mixed with the pre-formed RNP complex, and transferred to a single well of a 16-well cuvette. Electroporation was carried out using the program CA137. Immediately following nucleofection, 100 µL of culture medium was added, and cells were transferred to feeder cell-containing 6-well plates in epithelial cell-conditioned medium (prepared as per the *Fluorescence-activated cell sorting for clonal cell culture* section). Plates were incubated at 37°C and 5% CO₂ for 72 h.

To generate clonal cultures, epithelial cells were stained with anti-EPCAM-BV421 (BioLegend, 324220; 1:100 dilution) for 30 min at 4°C in FACS buffer, washed, and resuspended at 1 x 10⁶ cells/mL in cell culture medium. EPCAM⁺ single cells were sorted into individual wells of feeder cell-containing 96-well plates using a BD FACSAria Fusion or BD FACSAria III. Epithelial cell-conditioned medium was supplemented as above and added to each well (100 µL per well). An additional 50 µL of medium was added every 2-3 d until 7 d, after which full medium changes were performed. Confluent clones were sequentially expanded into 24-well and 6-well plates prior to downstream differentiation or analysis. In parallel, nucleofected epithelial cells were seeded directly onto transwells for differentiation in ALI culture and referred to as ‘bulk cultures’, as they contained mixed edited and unedited populations.

Genomic DNA was isolated from expanded clones using the AllPrep DNA/RNA Mini Kit (Qiagen, 80004) and quantified by NanoDrop (Thermo Fisher Scientific). Targeted genomic loci (*DNAH5* and *DNAI2*) were PCR-amplified using Q5 High-Fidelity DNA Polymerase (NEB) with primers listed in Table S8. 50 µL reactions contained 1× Q5 buffer, 0.02 U/µL polymerase, 50 ng genomic DNA, 200 µM dNTPs, and 0.5 µM of each primer.

Thermocycling was performed on a ProFlex thermocycler (Thermo Fisher Scientific) with an initial denaturation at 98°C for 30 s; 35 cycles of 98°C for 10 s, 50-72°C for 30 s, and 72°C for 30 s; followed by a final extension at 72°C for 2 min. A single amplification product was confirmed using 1% agarose gel, purified using ExoSAP-IT PCR Product Cleanup (Thermo Fisher Scientific, 75001.1.ML) and sent for Sanger sequencing (Source Bioscience).

#### Lentivirus production and cell transduction

The lentiviral vectors pHIV-Luc-ZsGreen (a gift from Bryan Welm; Addgene plasmid 39196, http://n2t.net/addgene:39196, RRID:Addgene_39196) or pCCL-CMV-EGFP (previously described (Lau et al., 2024; Yuan et al., 2015)) were produced in HEK293T cells by co-transfection with pMD2.G (a gift from Didier Trono; Addgene plasmid 12259, http://n2t.net/addgene:12259, RRID:Addgene_12259) and pCMVR8.74 (a gift from Didier Trono; Addgene plasmid 22036, http://n2t.net/addgene:22036, RRID:Addgene_22036) in jetPEI transfection reagent (Polyplus). Viral particles were collected 48 h after transfection and concentrated by combining the supernatants with PEGit concentrator (5×; System Biosciences, LV810A-1) overnight at 4°C. After centrifugation at 1,500 × g for 45 min, the supernatant was removed and the pellet was resuspended in 1/10th of the original volume of PBS. Concentrated supernatants were stored at −80°C until use. Virus was titrated by transduction of HEK293T cells and flow cytometry.

To spin-transduce primary epithelial cells, cells in suspension equivalent to 15,000-16,000 cells/cm^2^, epithelial medium containing 4 or 8 μg/mL polybrene and small molecules as indicated in Supplementary Fig. 9 were combined with virus particles at the indicated MOI. Cells were centrifuged at 920 × g for 1 h at 30°C and then incubated at 37°C, 5% CO_2_ for 5-7 h. After, medium was refreshed to remove the polybrene and the viral particles, and 3T3-J2 fibroblast feeder cells were added at 20,000 cells/cm^2^. Cells were expanded for 48 or 72 h before analysis by flow cytometry. After trypsinization in 0.05% trypsin/EDTA, cells were stained with either anti-EPCAM/CD326 (BV421; BioLegend, 324219) or anti-ITGA6/CD49f (PE-Dazzle; BioLegend, 313626) antibodies to allow exclusion of 3T3-J2 cells during analysis. Flow cytometry was performed on a BD LSRII cell analyzer (BD Biosciences). ZsGreen+ flow sorting was performed on a BD FACSMelody^TM^ Cell Sorter.

#### Copy number analysis

Low-pass whole-genome sequencing was performed by GENEWIZ (Azenta Life Sciences) with a mean coverage of 0.39-0.55x following alignment to hg38 and deduplication. FASTQ files were generated per sample using bcl2fastq v2.20. Sequencing adapters and low quality bases were trimmed using Trimmomatic 0.39 and cleaned reads were aligned to the Homo sapiens reference genome GRCh38 using Sentieon 202112.01 as part of the Azenta Life Sciences pipeline. Aligned BAM files, provided by GENEWIZ, were analyzed using CNAscreen v1.1.0 (Hynds, 2026) in R v4.5.3. Briefly, CNAscreen runs QDNAseq v1.46.0 (Scheinin et al., 2014)(Bioconductor 3.22). Reads with MAPQ < 37, unmapped reads, reads failing platform quality checks and reads flagged as duplicates were excluded, and the remaining reads were counted per read in 100 kb bins. Bins were filtered using QDNAseq’s residual and blacklist filters and excluded if mappability was <80% or proportion of characterized (non-N) bases was <99.9%, followed by GC/mappability correction, normalization, outlier smoothing and CBS segmentation (DNAcopy; α = 1e-10, undo.SD = 1) with a per-sample seed derived from a fixed base seed (170826) and the sample name.

Chromosome Y was excluded as all samples analyzed were XX. Genome-wide plots show log2 relative copy number per passing bin with segments overlaid and were plotted with ggplot2 v4.0.2.

#### In vivo experiments

Animal studies were approved by the University College London Biological Services Ethical Review Committee and licensed under UK Home Office regulations (project licence PP2060881). Transplantations were performed in male NOD *scid* gamma (NSG) mice.

For subcutaneous cell transplantations, basal cells expanded in FAD+Y+WS6 for 12-14 passages were separated from 3T3-J2 feeder cells by differential trypsinization. 2 x 10⁶ cells were resuspended in 400 μL growth factor–reduced Matrigel and kept on ice. Under 2-3% isoflurane anesthesia, 200 μL of the cell suspension was injected subcutaneously into each flank of NSG mice. After 12 weeks, mice were culled, and Matrigel plugs were excised and processed for histology.

For tracheal transplantation, basal cells transduced with the pHIV-Luc-ZsGreen lentiviral vector expanded in FAD+Y+WS6 for 2-6 passages, differentially trypsinized to remove 3T3-J2 feeder cells, and counted. Mice were pre-conditioned with 10 μL 2% (w/v) polidocanol in PBS by oropharyngeal delivery to disrupt the tracheal epithelium (Ma et al., 2025). 5-6 h later, 3.5-5 x 10⁶ cells were introduced in 25-30 μL epithelial cell culture medium by oropharyngeal instillation under 2-3% isofluorane anesthesia. Cell engraftment was monitored once weekly by supine IVIS imaging of luciferase activity under 2-3% isofluorane anesthesia. IVIS analysis was performed in Aura software; mean radiance within a 3x4 cm box centered on the neck is shown, the background measurement was obtained by averaging readings from the abdomen. At end points, mice were killed under anesthesia following IVIS imaging and tracheas were removed and opened longitudinally along the ventral surface. The tracheas were flattened under a microscopy slide in a small volume of PBS and ZsGreen was visualized using a Leica MC15 FC stereoscope (2-3.5× magnification). Brightness and contrast were adjusted using ImageJ. Tracheas were fixed in 4% PFA overnight at 4°C and paraffin embedded for histological analyses.

For serial transplantation, tracheas were digested in pronase (1.5 mg/mL) overnight at 4°C, the residual tracheal tissue was removed and cells digested with DNase I (0.5 mg/mL) before seeding on feeder layers in FAD+Y+WS6. Cells were expanded for two passages, flow sorted (BD FACSMelody) for ZsGreen+ and expanded for three further passages before transplant into naive animals (secondary recipients).

#### Data visualization and statistics

The schematic diagrams in Figures 1B, 3B, 4A, 5A, 6A, 7A/F, S9A, S9C and S11A were created using <u>BioRender.com</u>. Analyses and data visualization were performed in GraphPad Prism v10.6.0 and in RStudio [R v4.2.2] with the tidyverse [v2.0.0] packages: dplyr (https://dplyr.tidyverse.org/ v1.1.4), ggplot2 (https://ggplot2.tidyverse.org/ v3.5.1), ggpubr (https://rpkgs.datanovia.com/ggpubr/ v0.6.0), rstatix (https://rpkgs.datanovia.com/rstatix/ v0.7.2), tidyr (https://tidyr.tidyverse.org/ v1.3.1), stringr (https://stringr.tidyverse.org/ v1.5.1), tibble (https://tibble.tidyverse.org/ v3.2.1). FlowJo (v10.8.1) was used for flow cytometry data. Bar charts are reported as mean +/-standard deviation. All statistical tests are reported in figure legends. * = p<0.05, ** = p<0.01, *** = p<0.001, **** = p<0.0001.

## Data and code availability

Data used to support the claims of the paper are available in the manuscript text, tables, figures and supplementary materials. The low-pass WGS pipeline (CNAscreen v1.1.0), including documentation and a demonstration dataset derived from publicly available data, is archived at Zenodo (https://doi.org/10.5281/zenodo.24482398) and maintained on GitHub (https://github.com/EpiCENTR-Lab/CNAscreen). CiliaFreqMap is available via Zenodo (https://doi.org/10.5281/zenodo.17107574).

## Acknowledgements

We thank Kyle O’Sullivan and Rebecca Towns (University College London Biological Services, UK) for technical assistance, Ayad Eddaoudi and Panayiota Constandinou for help with flow cytometry and cell sorting, and Denise R. Bairros de Pilger (University College London and The Francis Crick Institute, UK) for access to an ECHO 550 liquid handler. We thank Stephen Hart (University College London, UK) and the Francis Crick Institute Cell Services Facility for providing the *BMI1*-transduced nasal cells and lung cancer cell lines used in Supplementary Fig. 1, respectively. Finally, we thank Sian Goldsworthy and UCL Genomics (University College London, UK) for assistance with low-pass whole-genome sequencing analysis.

## Author contributions

JCO, YI, SMJ and REH conceived the study. JCO, ASF and NAM performed cell culture experiments, analyzed data and visualized data. EKH performed PCD disease modeling studies using CRISPR-Cas9 editing, analyzed data and visualized data. DRP and REH performed and analyzed cell transplantation experiments. SKR assisted with the design of CRISPR-Cas9 editing experiments. JCO, MJR and CP performed biopsy flow sorting and colony counting. IR performed Ussing chamber experiments under the supervision of DLB. AS-I performed transmission electron microscopy (TEM) and RBD performed immunofluorescence staining and analysis of ciliated epithelial cells. AZS performed quantification of TEM. TEM and ciliary immunofluorescence analyses were supervised by RAH. MG performed HSC experiments under the supervision of WG. GB performed gastric organoid experiments under the supervision of GGG and PDC. JCO, ASF, BMA and ORP counted colony formation assays. JMO-G and AYK provided primary thymic epithelial cells. MJR, EFM and CRB performed nasal brush biopsies. DAM wrote a macro for and assisted with analysis of ciliary beat frequency. AFMD performed airway and lung organoid experiments. YI connected kenpaullone with WS6 and performed initial optimization experiments. REH performed lentiviral transduction experiments, analyzed low-pass WGS data and wrote the first manuscript draft. CO’C, SMJ and REH acquired funding for the project. All authors reviewed the manuscript and provided comments. JCO, ASF and REH revised the manuscript after co-author feedback.

---

**Supplementary Figure 1:**
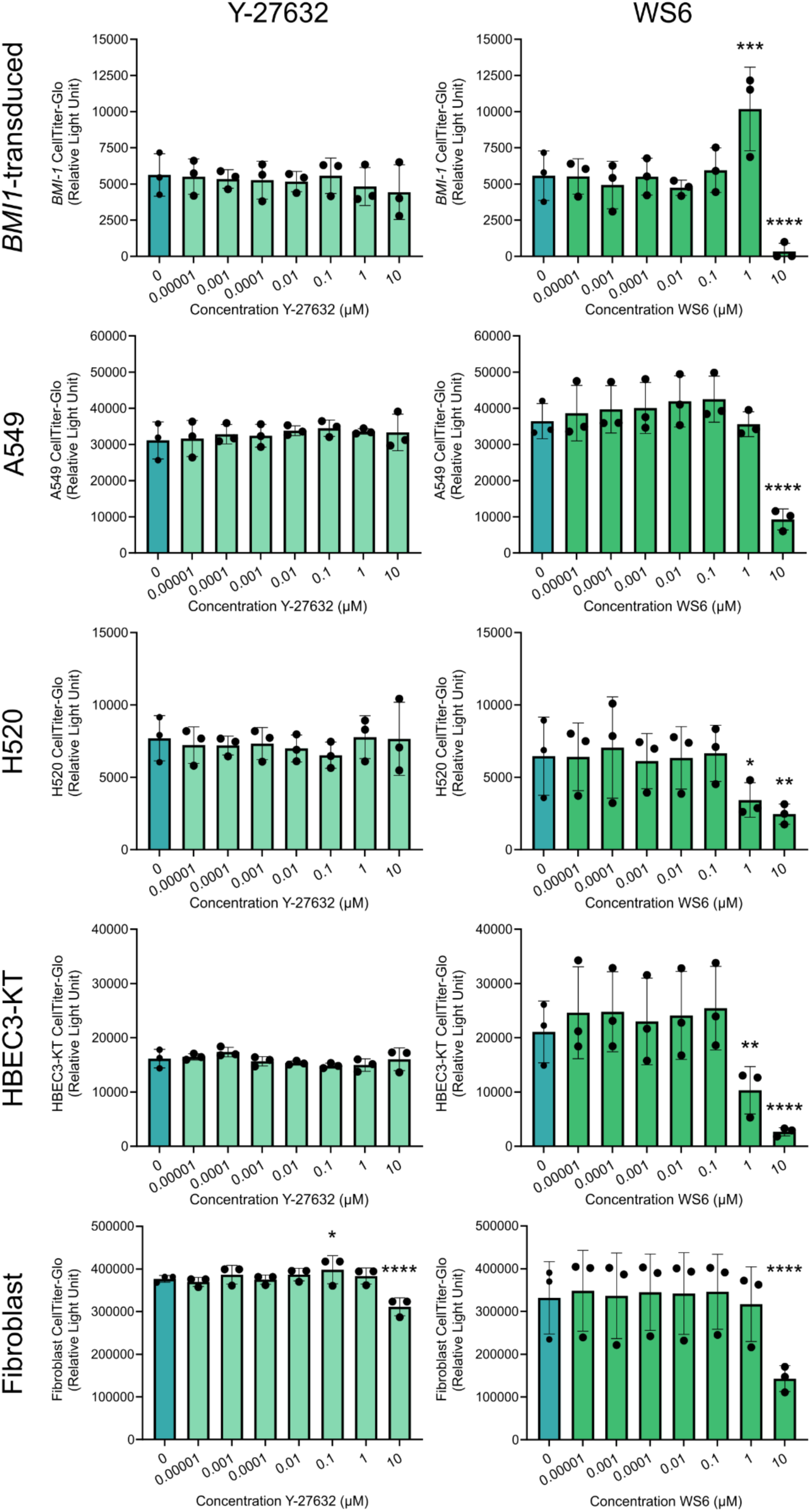
WS6 increases proliferation in *BMI1*-transduced cells but not other immortalized, cancer or fibroblast cell lines. Four-day CellTiter-Glo assays to assess the effect of varying concentrations of Y-27632 and WS6 on proliferation of epithelial cell lines (A549, lung adenocarcinoma; H520, lung squamous cell carcinoma; HBEC3-KT, *CDK4*-and *hTERT*-transduced bronchial epithelial basal cells; and *BMI1*-transduced nasal epithelial basal cells) and primary human lung fibroblast cell lines. Each data point represents the mean of technical triplicates for n = 3 experimental repeats except the primary human fibroblasts where n = 3 biological repeats. One-way ANOVA with Dunnett’s multiple comparison test vs DMSO only control wells.

**Supplementary Figure 2:**
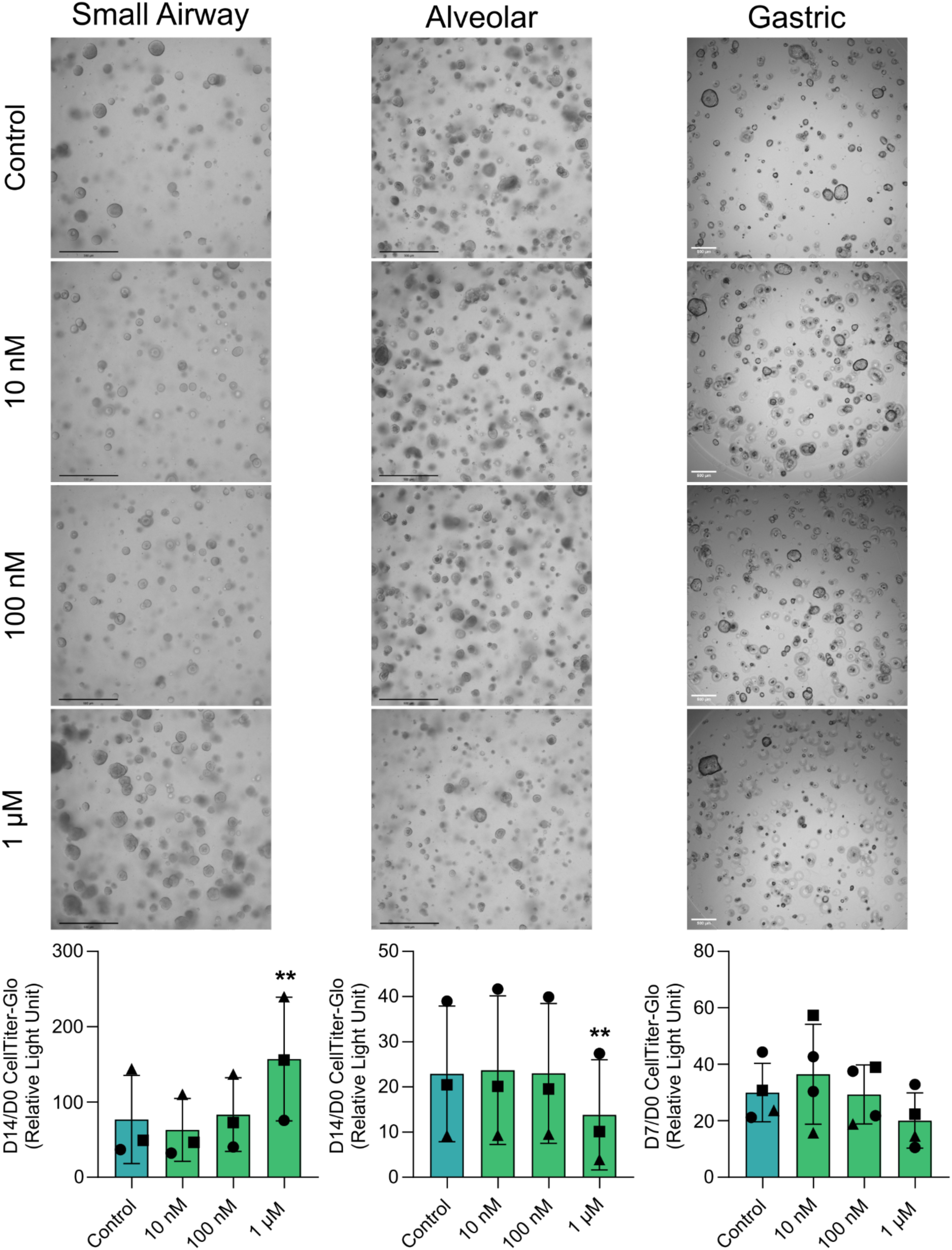
The effect of WS6 on primary human respiratory and gastric organoid cultures. Upper: Phase-contrast images of organoids treated with DMSO control, 10 nM, 100 nM, or 1 µM WS6. Scale bars = 500 µm. Lower: CellTiter-Glo endpoint/initial luminescence ratio to assess the effect of varying concentrations of WS6 on primary epithelial organoid cultures (airway and alveolar organoids: n = 3 biological replicates; gastric organoids: n = 2 donors, one with independent body and fundus organoid cultures, one with independent antrum and fundus organoid cultures; points represent the mean of triplicate wells per organoid culture). One-way ANOVA with Dunnett’s multiple comparison test vs DMSO control wells.

**Supplementary Figure 3:**
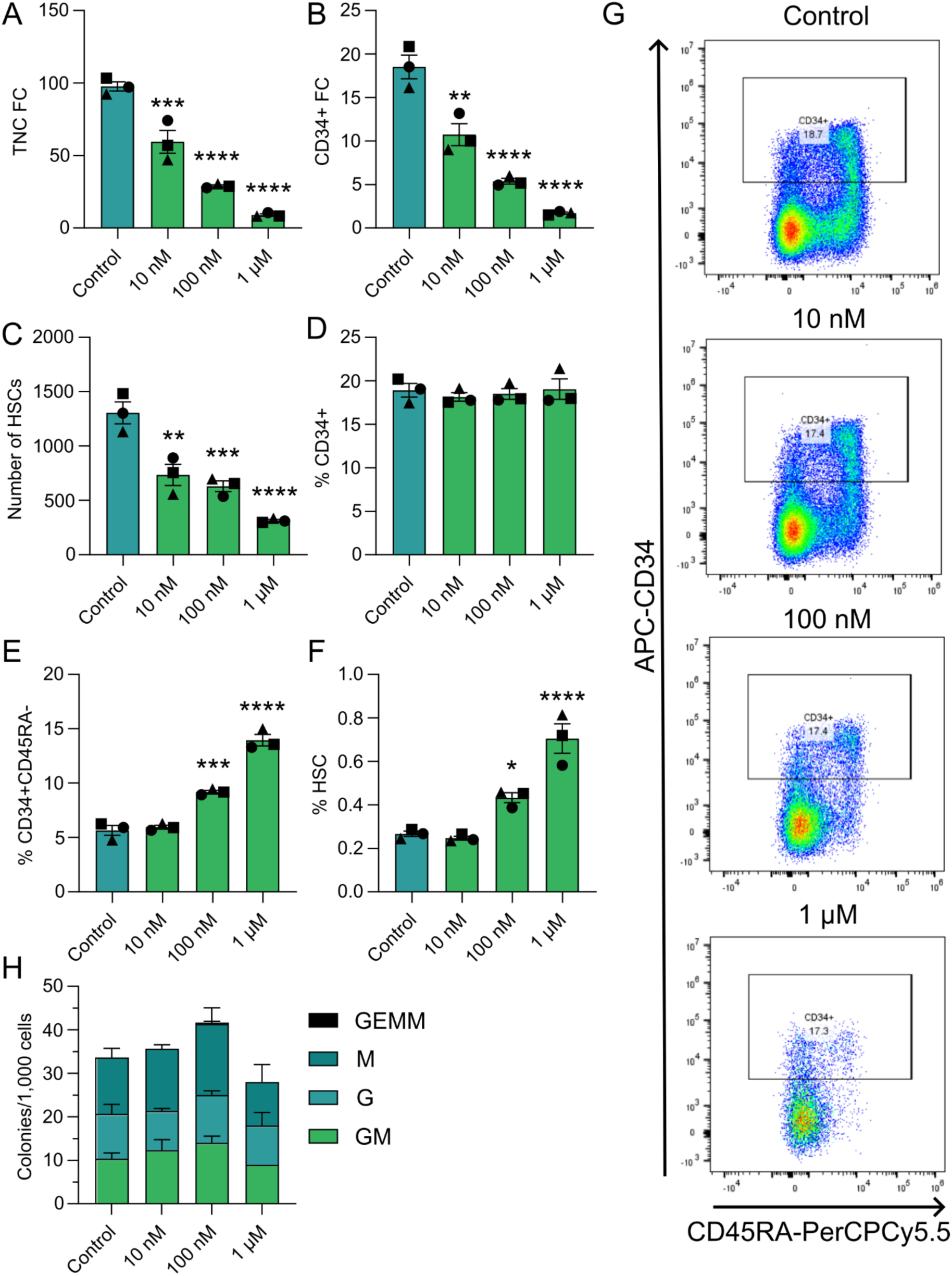
The effect of WS6 on primary human hematopoietic stem and progenitor cell cultures. 5,000 CD34^+^ cord blood stem and progenitor cells were cultured for 7 d in medium containing DMSO (control), 10 nM WS6, 100 nM WS6, or 1 µM WS6 before analysis by flow cytometry. A) Fold change in total nucleated cells relative to the number of cells seeded at day 0. B) Fold change in CD34+ cells relative to the number of cells seeded at day 0. C) Absolute number of hematopoietic stem cells (HSCs; CD34^+^, CD45RA^-^, CD90^+^, EPCR^+^). D) Proportion of CD34^+^ cells. E) Proportion of CD34+CD45RA^-^ cells. F) Proportion of HSCs. G) Representative flow cytometry plots showing CD34 and CD45RA expression. H) Colony forming unit (CFU) assay. CFU-macrophage (CFU-M), CFU-granulocyte (CFU-G), CFU-granulocyte macrophage (CFU-GM), and CFU-granulocyte erythrocyte macrophage megakaryocyte (CFU-GEMM). In all panels, one-way ANOVAs were performed with Dunnett’s multiple comparisons test comparing each condition to the control.

**Supplementary Figure 4:**
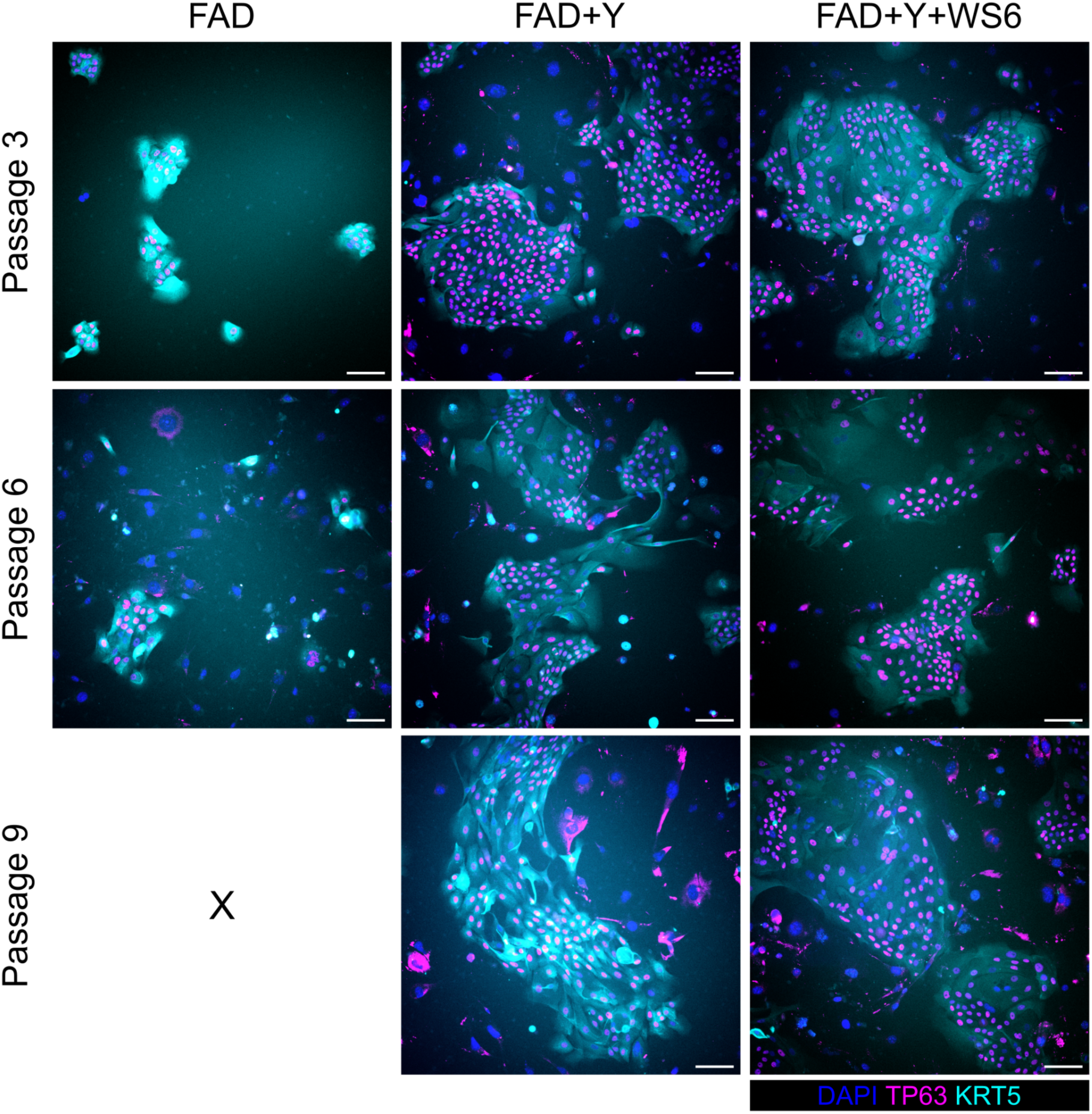
Epithelial basal cells retain KRT5 and TP63 expression during culture. Immunofluorescence staining of nasal basal cell cultures in FAD, FAD+Y-27632(Y) or FAD+Y+WS6 at the indicated passages (n = 1 donor displayed, n= 6 donors). Cells were fixed on day three of culture (TP63, magenta; KRT5, cyan; DAPI, blue). X represents passages at which imaging was not possible due to culture failure. Scale bars = 100 µm.

**Supplementary Figure 5:**
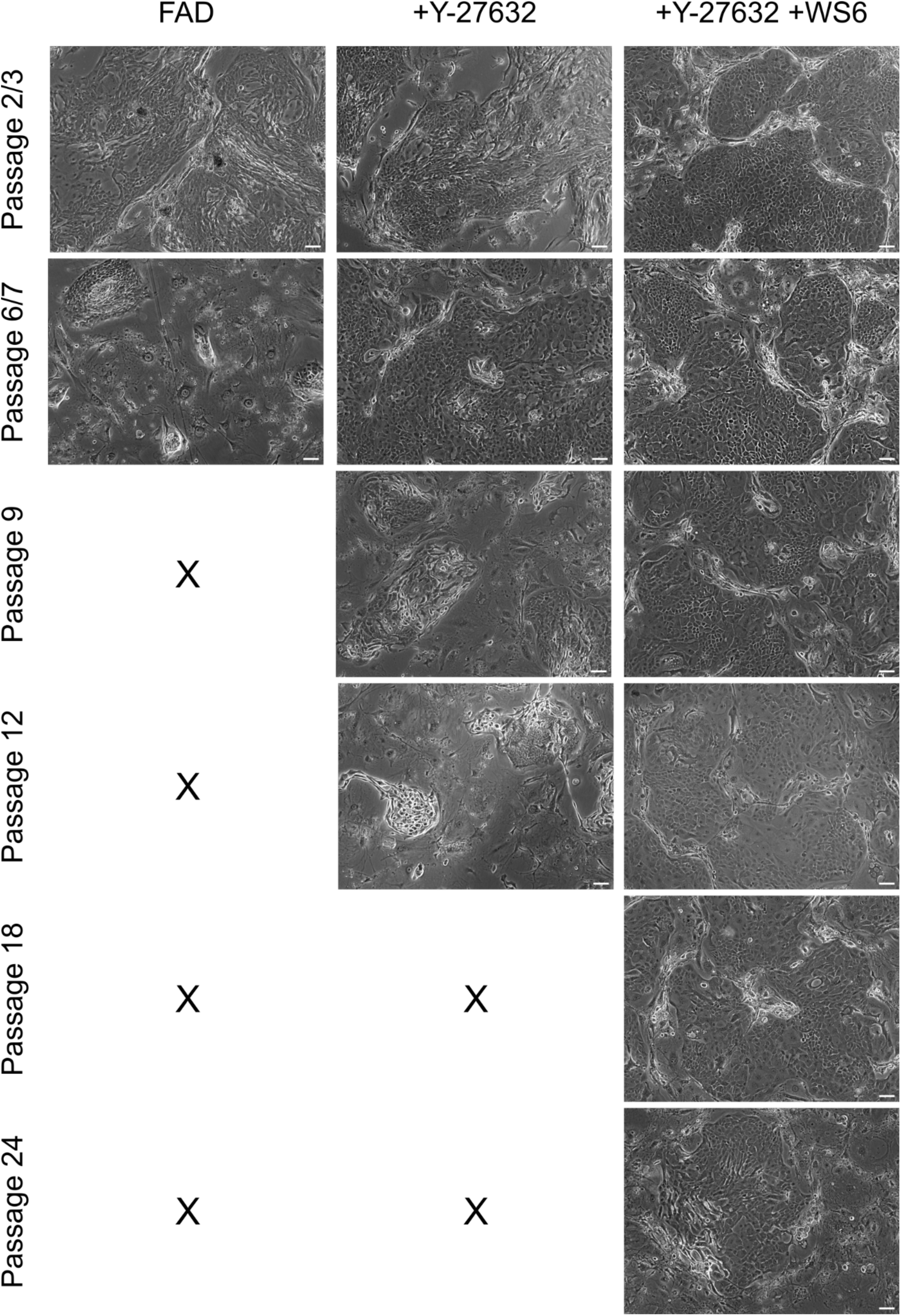
The morphology of epithelial basal cells cultured with WS6 remains consistent over long-term passage. Phase-contrast microscopy images show cultures in FAD, FAD+Y-27632(Y) or FAD+Y+WS6 at the indicated passages (n = 1 donor shown). X represents passages at which imaging was not possible due to culture failure. Scale bars = 100 µm.

**Supplementary Figure 6:**
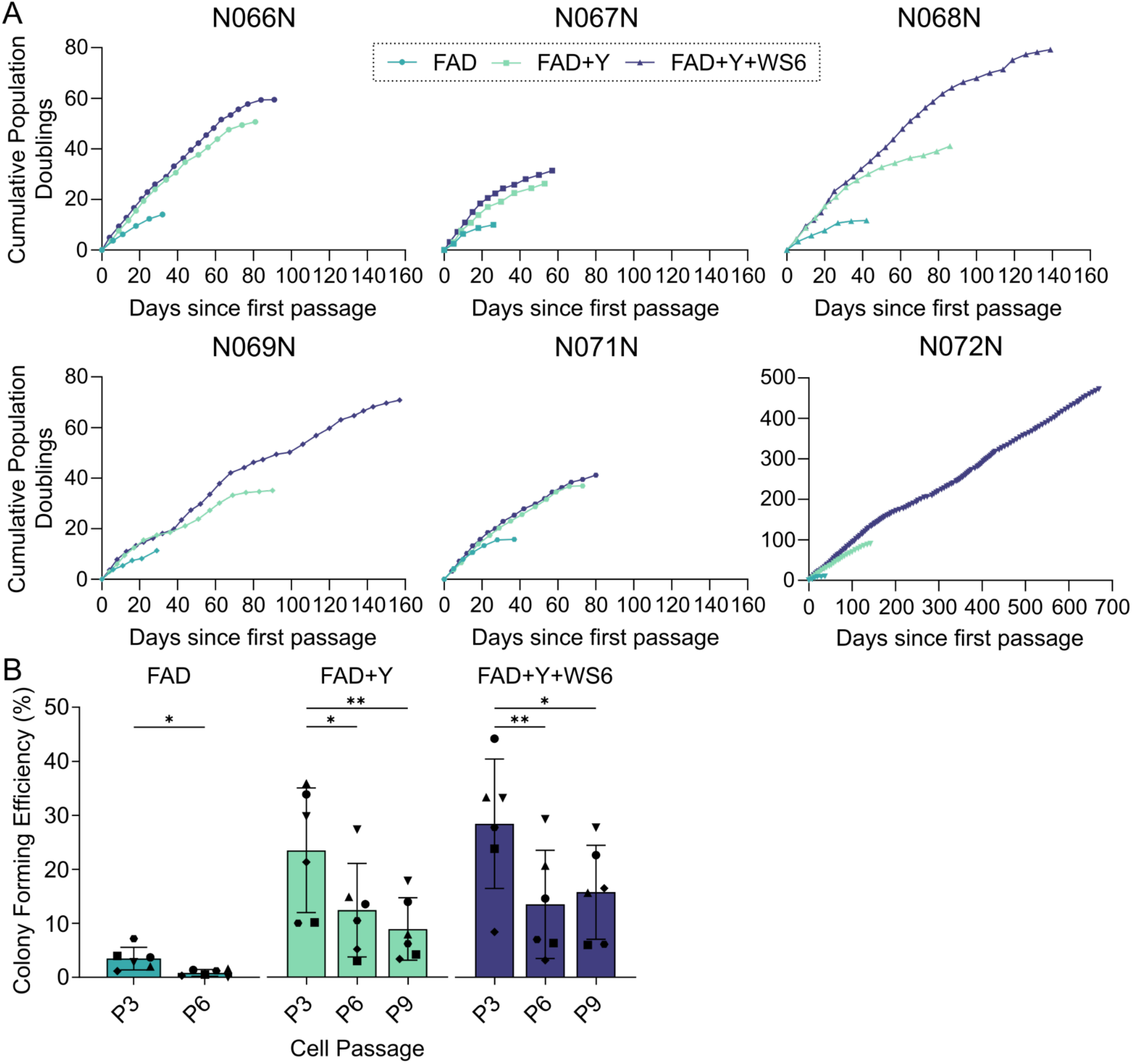
Epithelial basal cells in FAD+Y+WS6 have a longer cell culture lifetime. A) Cumulative population doublings of nasal epithelial basal cells cultured in FAD, FAD+Y or FAD+Y+WS6 from six individual nasal brush biopsies. Plot titles are donor IDs and N068N data is repeated from Fig 2E. B) Quantification of colony formation assays for donor-matched nasal epithelial basal cells cultured in FAD, FAD+Y or FAD+Y+WS6 at passages three, six, and nine (n = 6 donors). FAD condition = paired t-test, FAD+Y and FAD+Y+WS6 condition = one-way ANOVA with Tukey’s multiple comparison. Shapes indicate donors and data is repeated from Figure 2H.

**Supplementary Figure 7:**
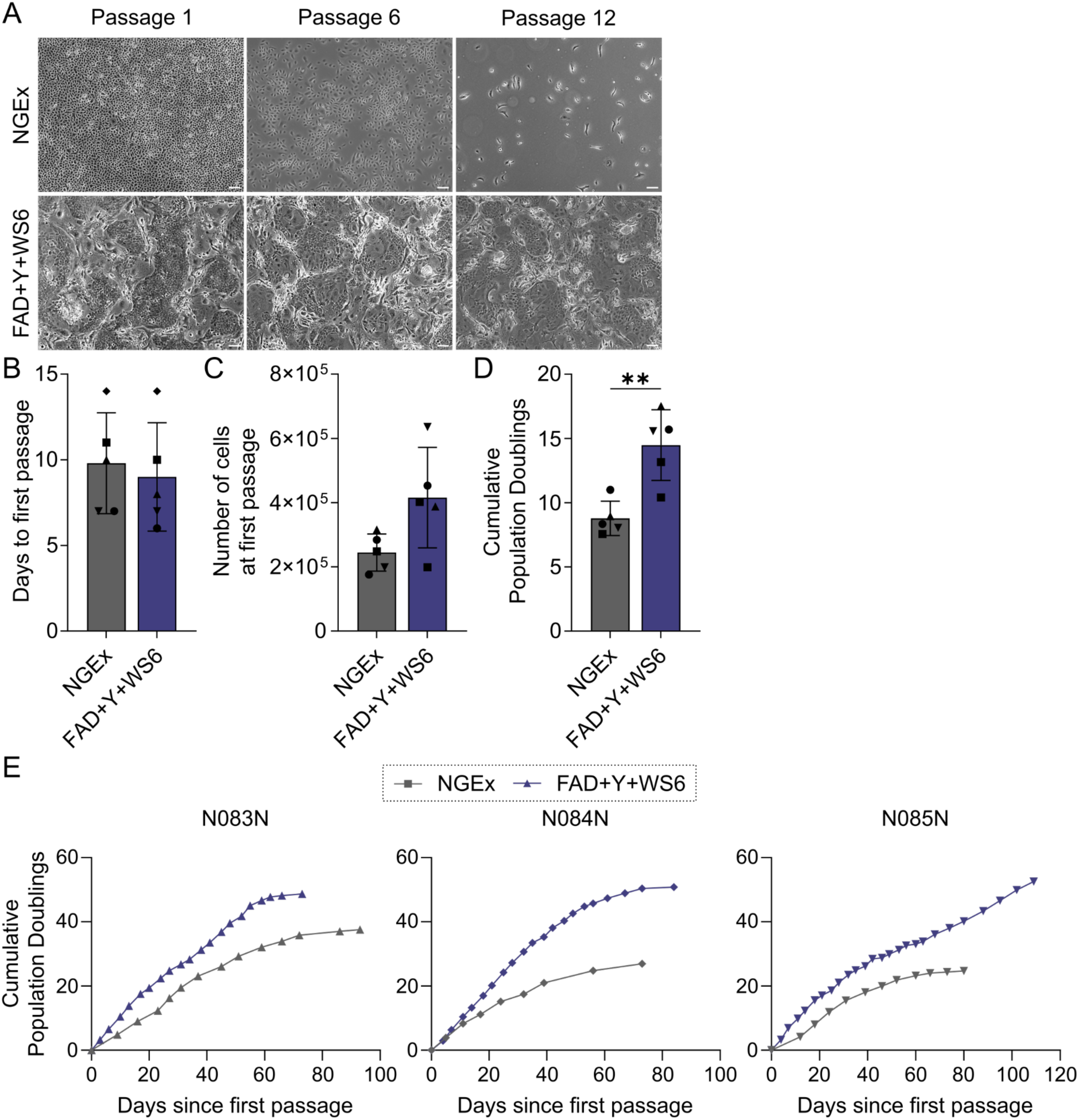
Increased expansion of nasal epithelial cells in FAD+Y+WS6 with 3T3-J2 co-culture compared to Pneumacult-NGEx. A) Representative phase-contrast microscopy images of nasal basal epithelial cultures in Pneumacult-NGEx (NGEx) and FAD+Y+WS6 at the indicated passages (n = 1 donor shown; passage 1 images on day of passage, passage 6 and 12 images are three days post seeding). Scale bars = 100 µm. B-C) Days to passage (B) and number of cells collected (C) at initial passaging of donor-matched nasal epithelial basal cells cultured in NGEx or FAD+Y+WS6 from a single brush biopsy from n = 5 donors. D) Cumulative population doublings for donor-matched nasal epithelial cells cultured in NGEx or FAD+Y+WS6 conditions on or after 21 days in cell culture (donor-matched values within one day of each other). B-D) Paired t-test. E) Cumulative population doublings of nasal epithelial basal cells cultured in NGEx or FAD+Y+WS6 conditions from three donors. Plot titles are donor IDs.

**Supplementary Figure 8:**
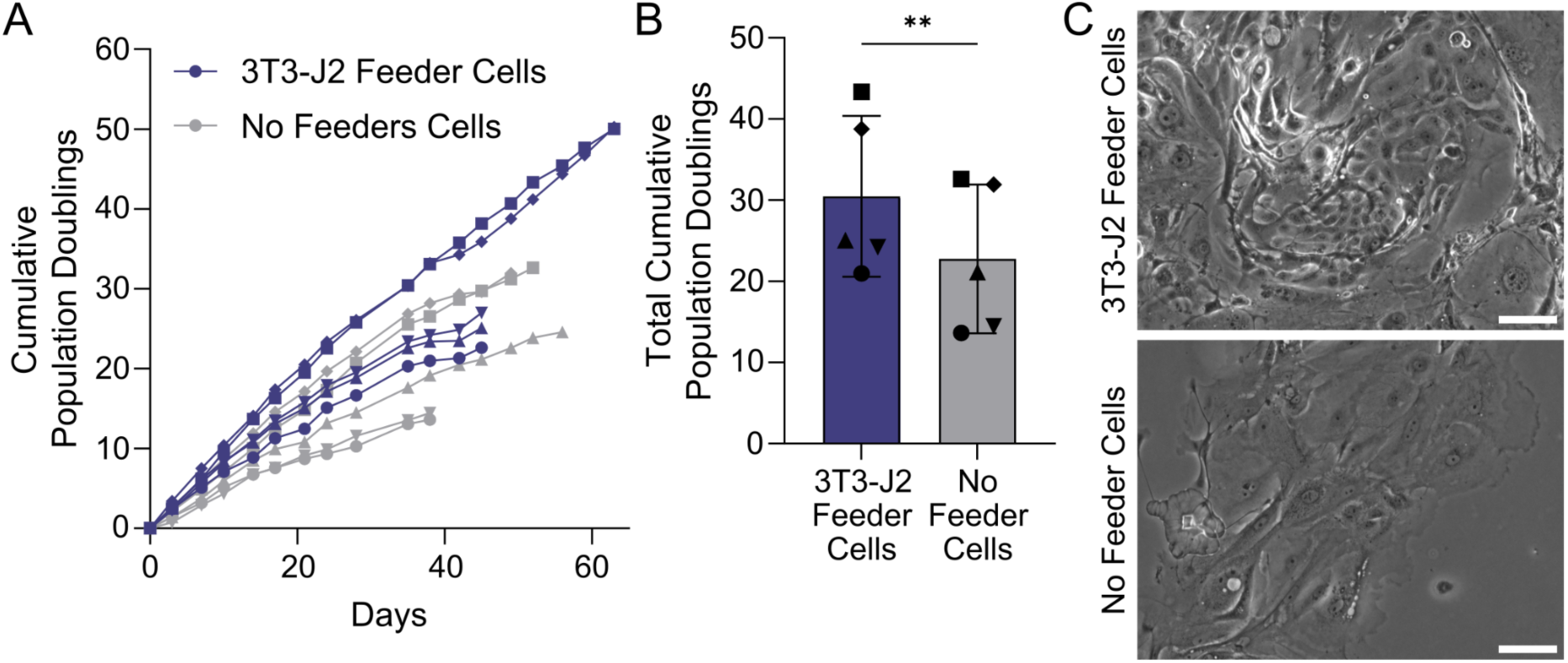
WS6 does not eliminate the use of 3T3-J2 fibroblast feeder cells. A) Cumulative population doublings of nasal epithelial basal cells cultured in FAD+Y+WS6 with 3T3-J2 feeder cells and with no 3T3-J2 feeder cells. Day 0 is the day of withdrawal or continuation of 3T3-J2 feeder cells after three passages in FAD+Y+WS6 with 3T3-J2 feeder cells (n = 5 donors, shape indicates donor). B) Cumulative population doublings from nasal epithelial cell cultures in FAD+Y+WS6 with 3T3-J2 feeder cells and with no 3T3-J2 feeder cells (n = 5 donors, shape indicates donor) where values are taken at the timepoint the first culture failed per donor. Paired t test. C) Phase-contrast microscopy of nasal epithelial cells cultured with and without 3T3-J2 feeder cells day 4 after passage 14. Scale bars = 100 µm.

**Supplementary Figure 9:**
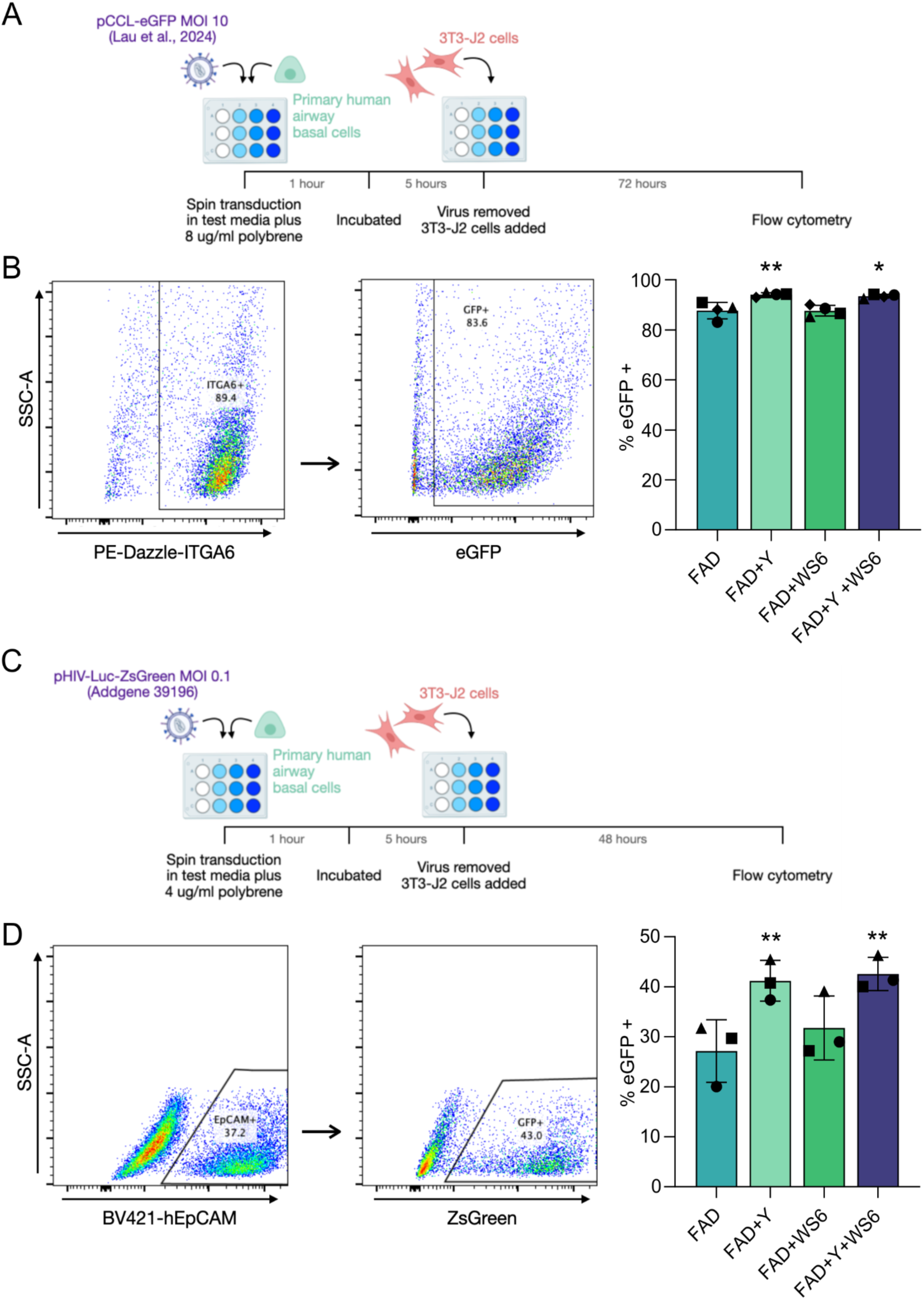
Y-27632 but not WS6 improves lentiviral transduction of primary epithelial cells. A) Bronchial basal cells previously cultured without Y-27632 (Y) or WS6 were spin transduced with a lentiviral vector carrying eGFP. B) Flow cytometry analysis of transduction efficiency in ITGA6-positive cells (exclusion of 3T3-J2 feeder cells) in control medium (FAD), FAD+Y, FAD+WS6 or FAD+Y+WS6. Each data point represents the mean of technical triplicates from one donor (n = 4 donors, one-way ANOVA with Dunnett’s multiple comparisons test comparing each condition to FAD). C) Bronchial and tracheal basal cells previously cultured without Y-27632 or WS6 were spin transduced with a lentiviral vector carrying ZsGreen. D) Flow cytometry analysis of transduction efficiency in EpCAM positive cells (exclusion of 3T3-J2 feeder cells) in control medium (FAD), FAD+Y, FAD+WS6 or FAD+Y+WS6. Each data point represents the mean of technical duplicates from one donor (n = 3 donors, one-way ANOVA with Dunnett’s multiple comparisons test comparing each condition to FAD).

**Supplementary Figure 10:**
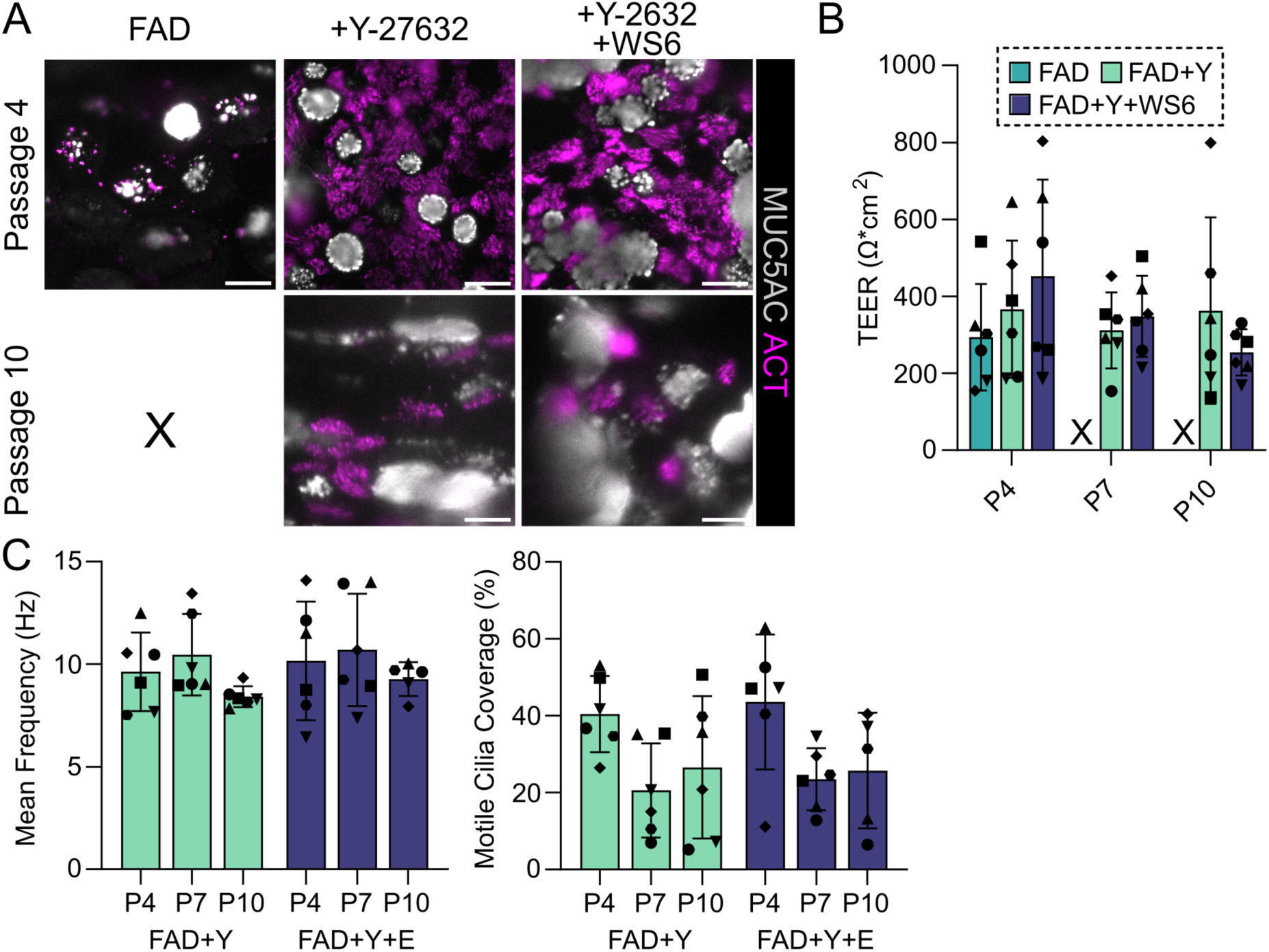
Nasal epithelial cells cultured in FAD+Y+WS6 retain the capacity for multipotent differentiation in air-liquid interface cultures. A) Immunofluorescence staining of passage four and ten ALI cultures of nasal epithelial basal cells from a single donor (n = 6 donors tested) from FAD (left), FAD+Y (center) or FAD+Y+WS6 (right) cultures. (ACT, magenta; MUC5AC, grey). Scale bars = 20 µm. B) Transepithelial electrical resistance (TEER) measurements of passage four, seven and ten air-liquid interface (ALI) cultures derived from nasal epithelial basal cells from FAD, FAD+Y-27632(Y) or FAD+Y+WS6 cultures. (n = 6 donors, passage 4 = one-way ANOVA with Tukey’s multiple comparison, passage 7 and 10 = paired t-test). C) Ciliary beat frequency analysis of differentiated ALI cultures from passage four, seven and ten cells previously expanded in FAD, FAD+Y or FAD+Y+WS6. Mean ciliary beat frequency (left) and motile cilia coverage (right). Each data point represents the average frequency of all moving cilia from five fields of view in three technical replicate wells (n = 6 donors; indicated by shape). Two-way ANOVA with Tukey’s multiple comparison, data for passage 4 ALI cell cultures are repeated from Figure 4E.

**Supplementary Figure 11:**
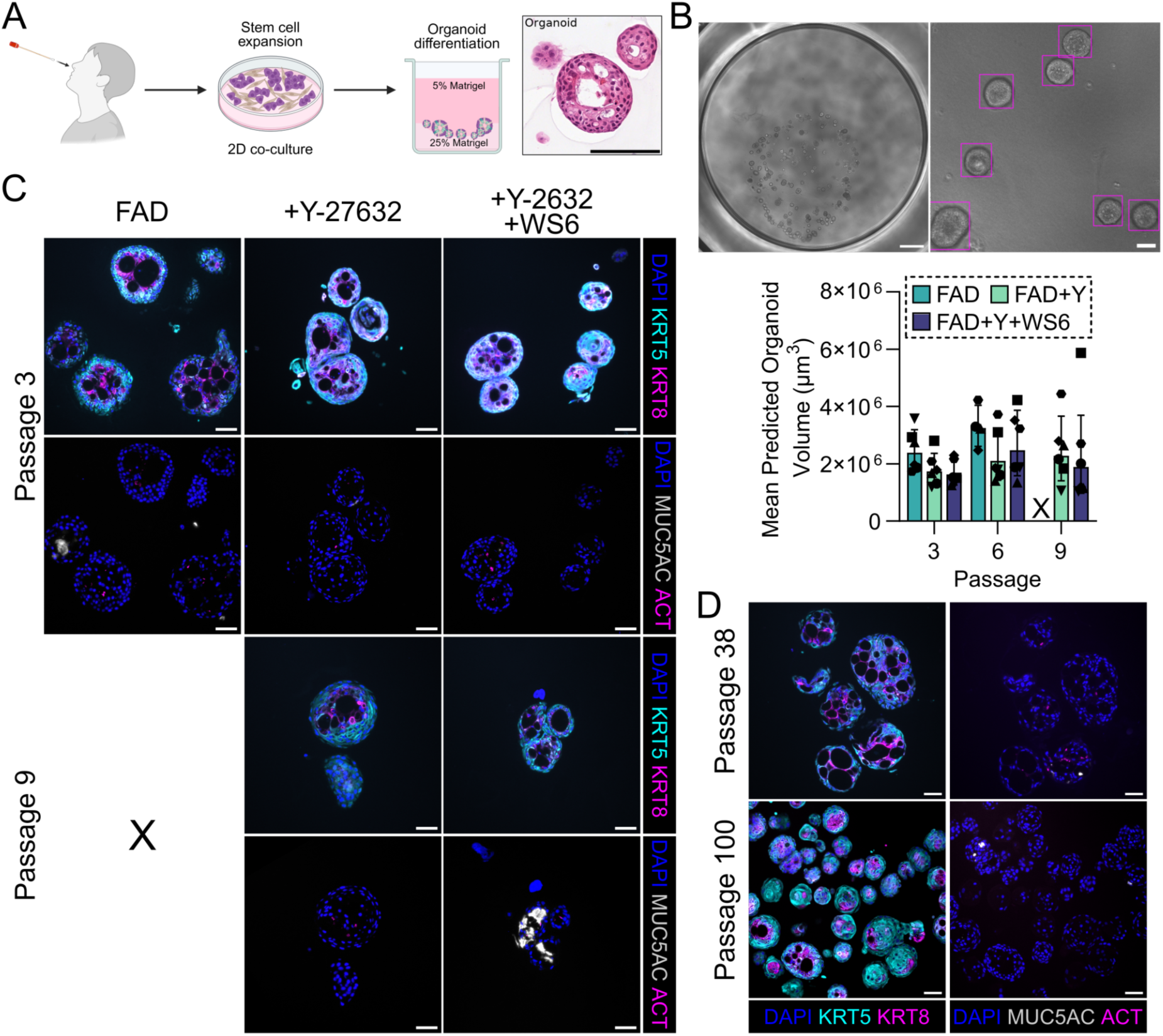
Nasal epithelial cells cultured in FAD+Y+WS6 retain the capacity for multipotent differentiation in organoid assays. A) Schematic diagram of nasal epithelial basal cell differentiation in organoid cultures with representative hematoxylin and eosin staining of formalin-fixed, paraffin-embedded organoid culture from a passage three FAD+Y+WS6 nasal epithelial basal cell culture. Scale bar = 100 µm. B) Whole-well phase-contrast images (upper left, scale bar = 1000 µm) at day 21/22 of organoid cultures derived from passage three, six or nine nasal epithelial basal cells from FAD, FAD+Y or FAD+Y+WS6 cultures. Images were analyzed using the OrgaQuant script to detect organoids (upper right, scale bar = 100 µm). Mean predicted organoid volume per condition is shown (lower; n = 6 donors, passage 3 = one-way ANOVA with Tukey’s multiple comparison, passage 6 and 9 = paired t-test). C) Immunofluorescence staining of passage three or passage nine organoid cultures of nasal epithelial basal cells from FAD (left), FAD+Y (center) or FAD+Y+WS6 (right) cultures. (upper: KRT5, cyan; KRT8, magenta; lower: ACT, magenta; MUC5AC, grey; DAPI, blue). Scale bars = 50 µm. D) Immunofluorescence staining of passage 38 and 100 organoid culture of nasal epithelial basal cells from FAD+Y+WS6 culture. (left: KRT5, cyan; KRT8, magenta; right: ACT, magenta; MUC5AC, grey; DAPI, blue). Scale bars = 50 µm. X represents passages at which assessment was not possible due to culture failure.

**Supplementary Figure 12:**
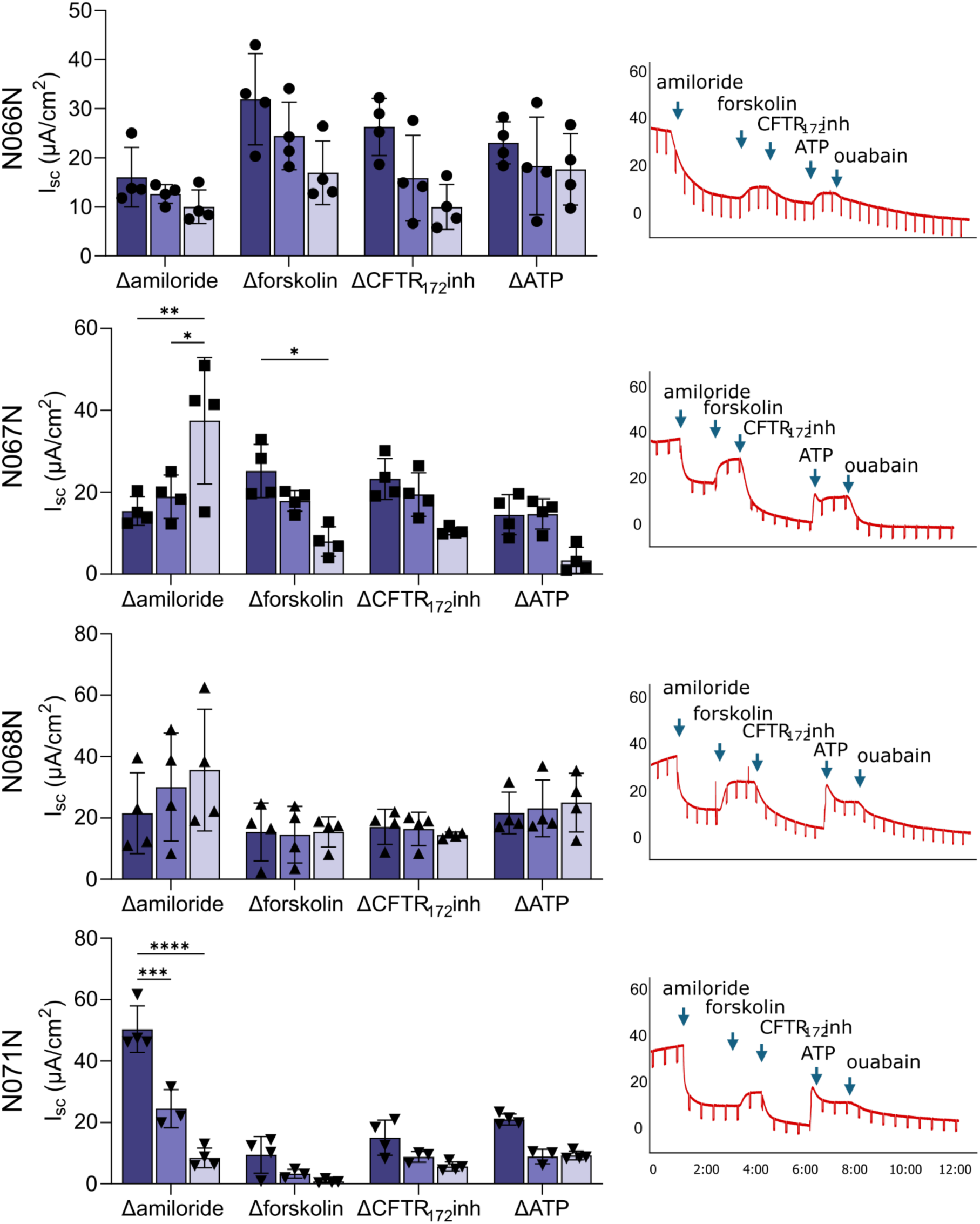
Ussing chamber analysis of ion transport in ALI cultures derived from FAD+Y+WS6 nasal cultures at passages four, seven and ten. Change in short-circuit current (I_sc_) of day 30 ALI cultures in response to amiloride (100 µM, apical), forskolin (10 µM, bilateral), CFTRinh-172 (10 µM, apical) and ATP (100 µM, apical), points represent individual well readings for each individual donor, n = 4 wells (left). Two-way ANOVA performed with Tukey’s multiple comparisons within the treatment group shown. Representative short circuit current (Isc) trace for each donor (right).

**Supplementary Figure 13:**
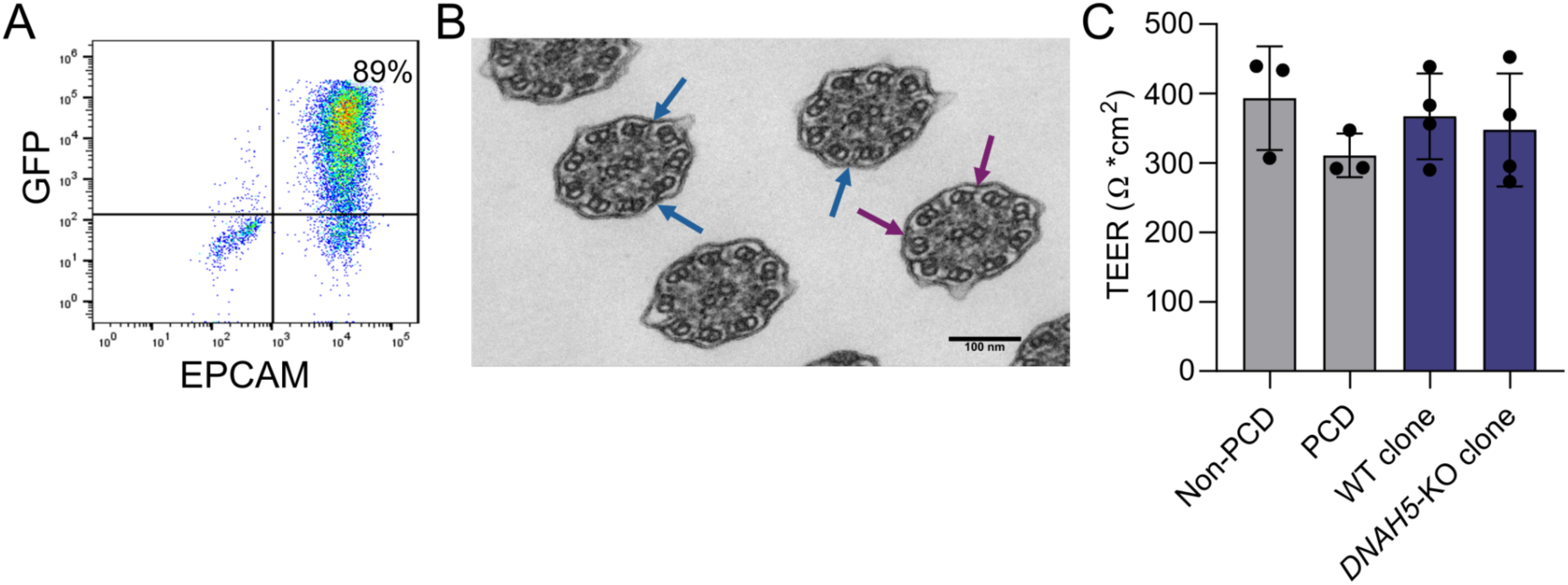
CRISPR-Cas9-mediated DNAH5-knockout in primary airway basal cells expanded in FAD+Y+WS6. A) Flow cytometry quantification of EpCAM⁺ epithelial cells expressing GFP following plasmid nucleofection, representing approximate nucleofection efficiency for single donor used for *DNAH5*-knockout (KO) experiment. B) Transmission electron microscopy of ciliary cross-section from the bulk population containing both *DNAH5*-KO and wild-type (WT) cells. Blue arrows indicate normal outer dynein arms (ODA) in WT cilia and purple indicate loss of ODA in *DNAH5*-KO. (40,000× magnification, scale bar = 100 nm). C) Transepithelial electrical resistance (TEER) measurements from air-liquid interface (ALI) cultures derived from *DNAH5*-KO clones, corresponding WT clones, non-primary ciliary dyskinesia (PCD) basal cells and PCD patient basal cells (n = 1 non-PCD and PCD patient donors with triplicate wells; n = 4 WT and *DNAH5*-KO clones from a single donor).

**Supplementary Figure 14:**
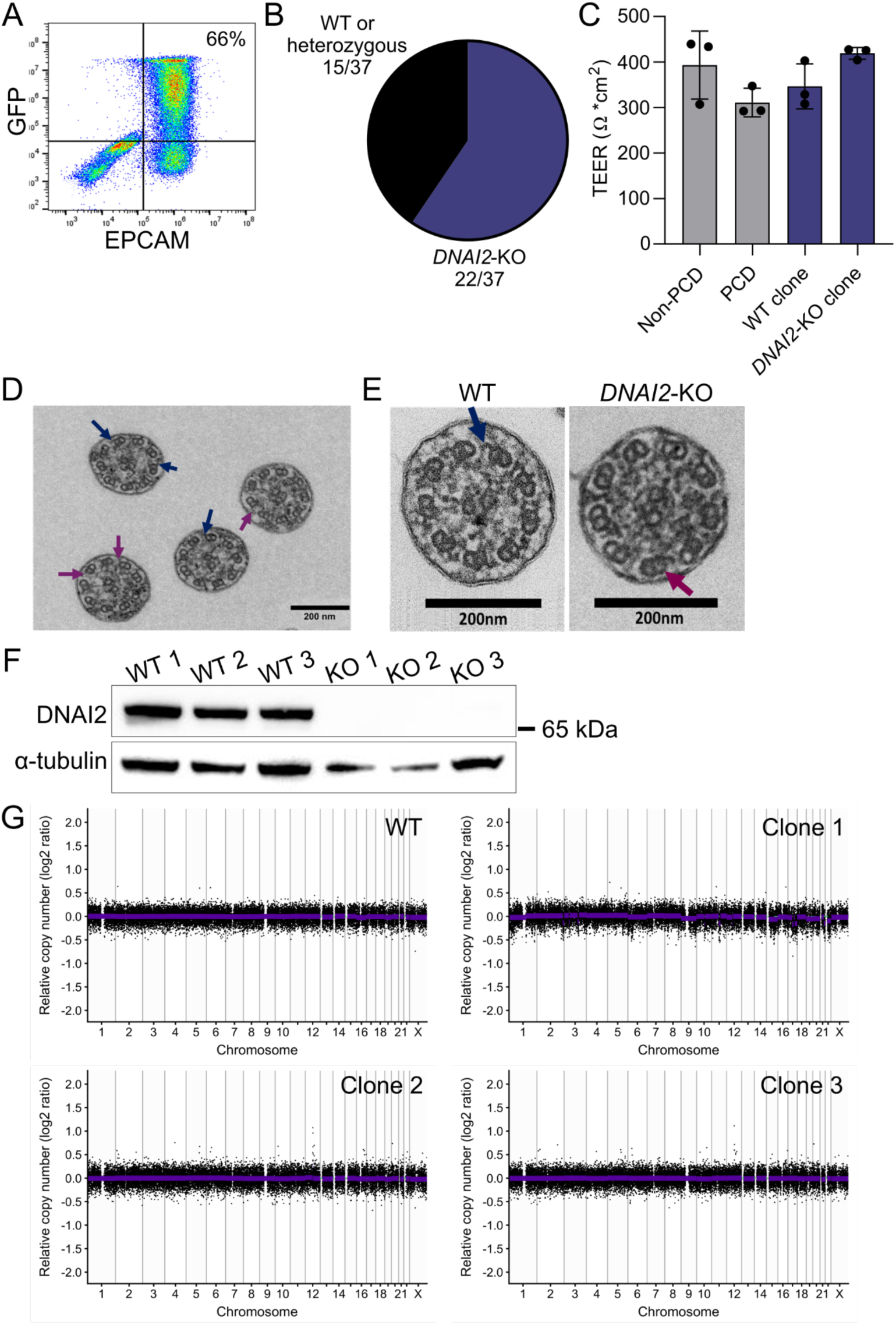
CRISPR-Cas9-mediated *DNAI2*-knockout in primary airway basal cells expanded in FAD+Y+WS6. A) Flow cytometry quantification of EpCAM⁺ epithelial cells expressing GFP following plasmid nucleofection, representing approximate nucleofection efficiency for single donor used for *DNAI2*-knockout (KO) experiment. B) Quantification of Sanger sequencing results showing the proportions of *DNAI2*-KO and wild-type (WT)/heterozygous clones. C) Transepithelial electrical resistance (TEER) measurements from air-liquid interface (ALI) cultures derived from *DNAI2*-KO clones, corresponding WT clones, non-primary ciliary dyskinesia (PCD) basal cells and PCD patient basal cells (n = 1 non-PCD and PCD patient donors with triplicate wells, n = 3 WT and *DNAI2*-KO clones; non-PCD and PCD control data are repeated from supplementary Figure 13C). D) Transmission electron microscopy (TEM) of ciliary cross-section from the bulk population containing both *DNAI2*-KO and unedited cells. Blue arrows indicate normal outer dynein arms (ODA) in WT cilia and purple indicate loss of ODA in DNAI2-KO in D and E. (40,000× magnification, scale bar = 200 nm). E) TEM images showing ciliary cross-sections from WT or *DNAI2*-KO clones differentiated in ALI cultures (n = 1 WT, n = 2 KO clones; scale bars = 200 nm). F) Western blot analysis of DNAI2 expression in WT clones (n = 3) and *DNAI2*-KO clones (n = 3). G) Copy number profiles from low-pass whole-genome sequencing of clonal nasal cell cultures expanded in FAD+Y-27632(Y)+WS6 (n = 1 WT clone, n = 3 *DNAI2*-KO clones). Points indicate unflagged bins and purple horizontal lines indicate copy number segments.

